# Assessing Mouse Kidney Parvovirus’ Ability to Confound Research by Examining its Effects on Renally Excreted Chemotherapeutics and its Impact on Pathologic Endpoints in the Adenine Model of Chronic Kidney Disease

**DOI:** 10.1101/2022.12.09.519764

**Authors:** Amanda C Ritter, Rodolfo J Ricart Arbona, Robert S Livingston, Sébastien Monette, Neil S Lipman

## Abstract

Mouse kidney parvovirus (MKPV) causes inclusion body nephropathy in severely immunocompromised mice and renal interstitial inflammation in immunocompetent mice. The purpose of this 2-part study was to determine the impact that MKPV may have on preclinical models as it relates to the pharmacokinetics of chemotherapeutics as well as its impact on the adenine diet model of chronic kidney disease. To assess the impact of MKPV on pharmacokinetics of 2 renally excreted chemotherapeutics commonly used in preclinical oncology studies, methotrexate and lenalidomide, blood and urine drug concentrations were measured in MKPV-infected or uninfected immunodeficient NOD.Cg-*Prkdc*^*scid*^*Il2rg*^*tm1Wjl*^/SzJ (NSG) and immunocompetent C57BL/6NCrl (B6) female mice. Differences in plasma pharmacokinetics were observed for methotrexate, but not for lenalidomide. Differences were most profound between uninfected NSG and B6 mice. The area under the curve (AUC) of methotrexate was 1.5-fold higher in uninfected NSG mice compared to infected NSG mice, 1.9-fold higher in infected B6 mice compared to uninfected B6 mice, and 4.3-fold higher in uninfected NSG mice compared to uninfected B6 mice. Renal clearance of both drugs was not impacted by MKPV infection but was generally lower in NSG mice. To assess the impact of MKPV on the adenine diet model of chronic kidney disease, MKPV-infected and uninfected B6 female mice were fed a 0.2% adenine diet and clinical and histopathologic features of disease were assessed over 8 weeks. Infection with MKPV did not have a significant impact on serum biomarkers of renal function such as BUN, creatinine, and SDMA; urine chemistry; or hemogram. However, infection did impact select histologic outcomes. MKPV-infected mice had significantly more foci of interstitial lymphoplasmacytic infiltrates than uninfected mice after 4 and 8 weeks of diet consumption, and significantly less interstitial fibrosis at week 8. Macrophage infiltrates and renal tubular injury, assessed using various immunohistochemical stains, were similar between groups. Together, these findings indicate that MKPV infection had minimal impact on the renal excretion of 2 chemotherapeutics and serum biomarkers of renal function. However, infection significantly impacted select histologic features of renal disease in the adenine diet model. While MKPV-free mice should be used in biomedical research, it is of the utmost importance in studies evaluating renal histology as an experimental outcome.

## Introduction

*Rodent chaphamaparvovirus 1* is a novel parvovirus species that is highly divergent from previously known parvoviruses of mice, and includes murine chapparvovirus (MuCPV) and mouse kidney parvovirus (MKPV)^43^. Although MKPV was only recently discovered in research mouse colonies, the histologic characteristics of the disease it causes, inclusion body nephropathy (IBN), have been noted by pathologists for over 40 years^4^. IBN is characterized by tubulointerstitial degenerative and inflammatory changes and intranuclear inclusions in tubular cells^4^. MKPV was discovered and demonstrated to be the causative agent of murine IBN in 2018^47^. MKPV is found frequently in research mouse colonies, with an estimated prevalence of approximately 10%^30^.

MKPV replicates predominantly in renal tubular epithelium (RTE), while extrarenal replication is minimal and restricted to very low copy numbers in the gastrointestinal mucosa^25; 30^. Once the virus has reached the kidney, the infection is persistent and results in continuous viral replication in RTE and shedding in urine for several months in both immunodeficient and immunocompetent mice^16; 25; 47^. Highly immunocompromised mice infected by MKPV are affected by a severe form of IBN, causing marked histopathologic lesions in kidneys, decreased renal function, and. morbidity and mortality^25; 47^ Although immunodeficient mice are most severely affected, immunocompetent mice also become infected and shed virus^16; 30; 47^. Infection in immunocompetent mice has been demonstrated to induce persistent interstitial nephritis, although clinical signs of disease have not been documented^16^. As this virus is known to persistently replicate in murine RTE cells and has the potential to induce pathologic changes without overt clinical illness, infection with this virus may impact renal function and related research^16; 25^.

Severely immunocompromised mice, including the NOD.Cg-*Prkdc*^*scid*^*Il2rg*^*tm1Wjl*^/SzJ (NSG) mouse, are routinely used in oncology research as they support the growth and maintenance of a broad array of human cancer cell lines and patient-derived xenografts (PDX)^24^. Therefore, the NSG mouse model often serves as a platform for testing the efficacy of emerging chemotherapeutics and combination therapies against various neoplasms, or as a patient surrogate for testing therapies in PDX^24; 35; 41^. Many chemotherapeutics depend on renal clearance, therefore the presence of a virus that impairs renal function may have substantial impact on research outcomes and may even impact patient care in the case of PDX-guided chemotherapy^22; 67^. Methotrexate and lenalidomide are chemotherapeutics commonly used to treat multiple cancers and autoimmune disorders in human patients. Both compounds are principally excreted via the renal tubules, albeit by different mechanisms. Sixty to 94% of methotrexate, an antifolate agent, is renally excreted^29^. It is actively excreted by numerous transporters in the proximal RTE, primarily the OAT3 and MRP4 transporters, and undergoes passive reabsorption in the distal renal tubules^22; 33^. Eighty-two percent of lenalidomide, a thalidomide analogue, is excreted primarily unchanged by kidneys using a P-glycoprotein transporter on the RTE^22^. It is not resorbed in the distal tubules nor does it bind to the OAT3 or MRP4 transporters used to excrete methotrexate^22; 29; 49^. As these compounds rely on the renal tubules for excretion, and MKPV primarily replicates in and causes damage to the renal tubules, we hypothesized that their renal excretion could be significantly impaired in mice infected with MKPV and may confound studies evaluating the use of chemotherapeutics in xenografted immunocompromised mice or immunocompetent mice engrafted with syngeneic tumors.

Murine models of renal physiology and pathophysiology allow researchers to address questions pertaining to various disease processes and help guide the development of novel therapies. The adenine diet model, which involves feeding mice a pelleted chow supplemented with 0.2% adenine, is a commonly utilized, well-established, murine model of chronic kidney disease (CKD)^23^. This diet induces a tubulointerstitial nephritis secondary to the metabolism of adenine by the enzyme xanthine dehydrogenase into 2,8-dihydroxyadenine (DHA) which precipitates forming crystals in the lumen of the proximal renal tubules and ascending limbs of the loops of Henle^28;59^. The resulting tubular injury leads to tubulointerstitial inflammation and fibrosis^28; 58^. Qualitative and quantitative assessment of renal tubular degeneration are outcomes routinely assessed in this model^2; 45; 50; 58^. The magnitude of fibrosis and decrease in renal function, assessed by measuring serum BUN, creatinine, and phosphorus, are commonly determined^2; 45; 50; 58^. Symmetric dimethylarginine (SDMA) has not been assessed in the adenine diet model, but as a sensitive marker of renal function in other species, it may have utility in future investigations of this and other models of renal disease in mice. As MKPV, like the dietary adenine model, primarily causes tubulointerstitial lesions, it is plausible that MKPV could also confound research outcomes in this model.

This study was conducted to ascertain whether infection with MKPV has the potential to confound research outcomes in commonly used murine models that depend on adequate baseline renal function. We hypothesized that infection with MKPV would decrease renal clearance of the chemotherapeutic agents methotrexate and lenalidomide, and that this effect would be exacerbated in an immunodeficient mouse strain. Additionally, we hypothesized that MKPV-infected mice would have significantly elevated biomarkers of renal injury such as serum BUN and creatinine, and that select histopathologic lesions would be more severe as compared to uninfected mice in the adenine diet model of CKD. This study also provided the opportunity to evaluate the novel renal serum biomarker, SDMA, in the adenine diet model as this marker has been shown to have advantages over serum creatinine as an indicator of renal function in other species, but its utility in mice has not been assessed^20; 40^.

## Materials and Methods

### Animals

Six to 8-week-old female C57BL/6NCrl (B6; Charles River Laboratories [CRL], Senneville, Quebec) and NOD.Cg-*Prkdc*^*scid*^ *Il2rg*^*tm1Wjl*^/SzJ (NSG; The Jackson Laboratory, Bar Harbor, ME) were used. All mice were free of mouse hepatitis virus, Sendai virus, mouse parvovirus, minute virus of mice, murine norovirus, pneumonia virus of mice, Theiler meningoencephalitis virus, epizootic diarrhea of infant mice (mouse rotavirus), ectromelia virus, reovirus type 3, lymphocytic choriomeningitis virus, K virus, mouse adenovirus 1 and 2, polyoma virus, murine cytomegalovirus, mouse thymic virus, hantavirus, mouse kidney parvovirus, *Mycoplasma pulmonis, Citrobacter rodentium, Salmonella* spp., *Filobacterium rodentium, Clostridium piliforme, Corynebacterium bovis*, fur mites (*Myobia musculi, Myocoptes musculinis*, and *Radfordia affinis*), pinworms (*Syphacia* spp. and *Aspiculuris* spp.), and *Encephalitozoon cuniculi* when the studies were initiated, as determined by testing naïve outbred Swiss Webster (Tac:SW) mice exposed repetitively to soiled bedding from cages housing mice in the colony.

### Husbandry and Housing

Mice were maintained in autoclaved individually ventilated polysulfone cages with stainless-steel wire-bar lids and filter tops (no. 19, Thoren Caging Systems, Inc., Hazelton, PA) on autoclaved aspen chip bedding (PWI Industries, Quebec, Canada) at a density of no greater than 4 mice per cage. Each cage was provided with a Glatfelter paper bag containing 6 g of crinkled paper strips (EnviroPak, WF Fisher and Son, Branchburg, NJ) for enrichment. Mice were fed a natural ingredient, closed source, autoclavable diet (5KA1, LabDiet, Richmond, VA) or a purified diet containing 0.2% adenine (TD.140290, Envigo, Madison, WI) ad libitum. All animals were provided autoclaved reverse osmosis acidified (pH 2.5 to 2.8 with hydrochloric acid) water in polyphenylsulfone bottles with stainless-steel caps and sipper tubes (Techniplast, West Chester, PA) ad libitum. Cages were autoclaved (Century SLH Scientific, Steris, Mentor, OHOH) with a pulsed vacuum cycle of 4 pulses at a maximum pressure of 12.0 psig, with sterilization temperature of 250.5°F (121.39°C) for 20 minutes, and a 10.0 inHg vacuum dry. Sterilization was confirmed by color change of temperature sensitive autoclave tape (Medline, Mundelein, IL) and review of the post-cycle chamber conditions. Water bottles were autoclaved at a temperature of 250.0°F (121.11°C) for 45 minutes with a purge time of 10 minutes. Cages were changed every 7 days within a class II, type A2 biological safety cabinet (LabGard S602-500, Nuaire, Plymouth, MN). The rooms were maintained on a 12:12-h light:dark cycle, relative humidity of 30% to 70%, and room temperature of 72 ± 2 °F (22.2 ± 1.1 °C). The animal care and use program at Memorial Sloan Kettering Cancer Center (MSK) is accredited by AAALAC International, and all animals are maintained in accordance to the recommendations provided in the Guide^21^. All animal use described in this investigation was approved by MSK’s IACUC in agreement with AALAS’ position statements on the Humane Care and Use of Laboratory Animals and Alleviating Pain and Distress in Laboratory Animals.

### MKPV Infection

Viral stock (MKPV substrain MSK-WCM2015-3781-1-2) was created by thawing and homogenizing frozen kidney obtained from naturally infected NSG mice with histologically and PCR confirmed inclusion body nephritis (IBN) caused by MKPV infection^30^. The homogenate was resuspended in 1X PBS, passed through a sterile 0.22 μm filter (MilliporeSigma, Burlington, MA), and then centrifuged at 626 x g for 5 minutes. Supernatant was collected, aliquoted, and stored at -80°F. This viral stock was thawed once, diluted to 1:100 using sterile PBS, aliquoted into individual portions, and stored at -80°F until ready for use. At the time of inoculation, individual aliquots were thawed and administered to each mouse via oral gavage (200 μL viral stock) and intranasally (25 μL of viral stock per nostril; 50 μL total volume). Frozen kidney samples from mice in all sham-inoculated groups were submitted for PCR to ensure animals remained MKPV-negative for the duration of the study.

### MKPV PCR

Samples were analyzed using a validated proprietary PCR assay (IDEXX Laboratories, Columbia, MO). Total nucleic acids were extracted from kidney or urine using a commercial kit (NucleoMag Vet, Machery-Nagel, Düren, Germany). The real-time PCR assay targeted the gene coding for the MKPV NS1 protein (GenBank NC040843.1) using a hydrolysis probe (Integrated DNA Technologies, Coralville, IA). The assay has an analytical sensitivity of between 1 and 10 template copies per PCR reaction. Analysis and amplification of the product were performed using standard primer and probe concentrations with a commercially available master mix (LC480 ProbesMaster, Roche Applied Sciences, Indianapolis, IN) on a commercially available real-time PCR platform (LightCycler 480, Roche Applied Sciences). The copy number estimate of MKPV DNA in each PCR test was calculated by plotting the real-time crossing point values from the MKPV PCR assay on a standard curve generated by testing log-fold dilutions of a known copy number positive control.

### Pharmacokinetics Study

#### Experimental Design

Adult female NOD.Cg-*Prkdc*^*scid*^ *Il2rg*^*tm1Wjl*^/SzJ (NSG) and C57BL/6NCrl (B6) mice (6-8 weeks of age; n=48 per strain) were used. Mice were randomly assigned by a random number generator (Randomizer.org) to 1 of 2 treatment groups: inoculation with MKPV (n=24 mice per strain) or sham inoculation with an equivalent volume of sterile phosphate buffered saline (PBS) (n=24 mice per strain). Ten to 11 weeks following inoculation, urine was collected from each mouse and assayed for MKPV by PCR. Fourteen weeks after inoculation, mice (n=12 per group) were administered a single intravenous bolus of either 5 mg/kg lenalidomide or 70 mg/kg methotrexate by tail vein. Blood was collected post-lenalidomide administration at 5, 15, 30, 60, 120, and 240, 360, and 480 minutes, and post-methotrexate administration at 5, 15, 30, 45, 60, 90, 120, and 240 minutes. Blood was collected from each mouse twice, a survival collection by submental puncture and a terminal collection performed by retro-orbital puncture under isoflurane anesthesia, followed by euthanasia by CO_2_ asphyxiation. Blood samples were collected from 3 mice at each time point and assayed for lenalidomide or methotrexate. Between the survival and terminal blood collection, mice were housed individually in cages with hydrophobic sand (Lab Sand, Coastline Global Inc, West Chester, PA) to facilitate urine collection. All urine excreted between the survival and terminal timepoint was collected for each mouse, the volume measured, and assayed for lenalidomide or methotrexate. After euthanasia, kidneys were removed, the cranial pole of the left kidney was frozen at -80°C and retained for MKPV PCR, and the remaining renal tissue was preserved in 10% neutral buffered formalin for histopathology. Additionally, blood collected at the terminal time point was analyzed for serum BUN, creatinine, and phosphorus.

#### Compound Preparation

Lenalidomide (SML2283, Sigma-Aldrich, St. Louis, MO) was dissolved in dimethyl sulfoxide (DMSO) then diluted with sterile PBS containing 1% hydrochloric acid (HCl) to achieve a 1 mg/mL lenalidomide solution. Methotrexate 25 mg/mL (NDC 16729-277-30, Accord Healthcare, Inc., Durham, NC) was diluted with sterile saline to 10 mg/mL. All solutions were diluted less than 2 hours prior to use and sterile filtered using a 0.22 μm filter (MilliporeSigma).

#### Pharmacokinetic Quantification

Whole blood was collected into tubes containing anti-coagulant (MiniCollect K2E K2EDTA, Greiner Bio-One, Monroe, NC) and centrifuged at 626 x g at 4°C for 10 minutes. Plasma was decanted and placed in a preservative-free collection tube (Microcentrifuge tube, Globe Scientific, Mahwah, NJ) and stored at -80°C until analysis. Urine was collected in preservative-free collection tubes (0.2mL PCR Tube, Eppendorf, Hamburg, Germany) and stored at -20°C until analysis. LC-MS analysis of thawed plasma and urine samples was performed on a triple quadrupole mass spectrometer (LCMS-8030 with LC-20AD LC pumps, Shimadzu, Kyoto, Japan). Separation was performed using an HPLC column (Zorbax Eclipse XDB-C18 [2.1 × 50 mm, 3.5 μm], Agilent, Santa Clara, CA). Gradient separation was performed using 0.1% formic acid in water (aqueous, A) and 0.1% formic acid in acetonitrile (organic, B) via the following gradient (flow rate – 0.5 mL/min): 2-30% organic from 0-3 minutes, followed by a two-minute hold at 100% B and a two-minute return to starting conditions (2% B). Mass spectrometric detection was performed using multiple reaction monitoring (positive mode) of the following optimized transitions for Lenalidomide and Methotrexate: Lenalidomide: 259.9 → 149.05; 259.69 → 187.10; Methotrexate: 454.9 → 308.05; 454.9 → 175.10.

#### Pharmacokinetic Parameter Calculations

A non-compartmental pharmacokinetic (PK) analysis was performed with software (PKanalix version 2020R1, Lixoft, Orsay, France), using the linear-log trapezoidal rule for calculation of the area under the concentration-time curve (AUC_last_). The PK parameters determined were the area under the concentration-time curve from 0 to the last observation (AUC_last_), back-extrapolated plasma drug concentration at time zero (C0), systemic clearance (Cl), and elimination half-life (t1/2). Data points below the limit of quantification (LOQ) were replaced by the LLOQ (lower limit of quantification) divided by 2. Renal clearance (CL_R_) was calculated by the Ae_t1–t2_/AUC_t1–t2_ ratio where Ae_t1-t2_ was calculated by the summation of drug excretion (urine drug concentration multiplied by urine volume) during the urine collection period t1(survival blood collection) to t2 (terminal blood collection)^64^. Urine collection intervals for methotrexate were: 5-60 minutes, 15-90 minutes, 30-120 minutes, and 45-240 minutes post-administration; urine collection intervals for lenalidomide were: 5-120 minutes, 15-240 minutes, 30-360 minutes, and 60-480 minutes post-administration. Mean (standard deviation), geometric mean (95% confidence interval [CI]), and coefficient of variation (%CV) were calculated for each plasma pharmacokinetic parameter. Mean and standard deviation were determined for calculated renal clearances.

#### Clinical Pathology

For serum chemistry analysis, clotted blood collected in serum separator tubes (MiniCollect 0.8mL CAT Serum Separator, Greiner Bio-One, Monroe, NC) was centrifuged at 3913 x g for 10 minutes, the serum decanted and stored at -80°C until analysis. Thawed serum samples were processed using an automated analyzer (AU680 Chemistry Analyzer, Beckman Coulter, Brea, CA). The serum concentration of blood urea nitrogen (BUN), creatinine (CREA), and phosphorus (P) were determined.

#### Histopathology

Following euthanasia by CO_2_ asphyxiation, a complete post-mortem gross examination was performed, gross lesions were recorded, and kidneys were fixed in 10% neutral buffered formalin. After 24-48 hours of fixation, kidneys were trimmed into two longitudinal and one transverse sections per mouse, processed in alcohol and xylene, paraffin-embedded, sectioned at 5 microns and stained with hematoxylin and eosin (H&E). Renal histopathology was semi-quantitatively scored as previously described^16^. In brief, slides were reviewed for tubular changes and inflammatory infiltrates, and these features were graded on a scale from 0 (within normal limits) to 4 (marked inflammatory infiltration or tubular changes). Intranuclear inclusions were noted to be absent or present.

#### RNA in situ hybridization (ISH)

Renal tissue from animals which were inoculated with, but tested negative for MKPV via PCR, were evaluated for the presence of MKPV RNA by in situ hybridization (ISH) using a target probe designed and validated to detect both viral capsid (VP1) and non-structural (NS1) regions of MKPV^47^. Slides were stained on an automated stainer (Leica Bond RX, Leica Biosystems) with RNAscope 2.5 LS Reagent Kit-Red (322150, Advanced Cell Diagnostics, Newark, CA) and Bond Polymer Refine Red Detection (DS9390, Leica Biosystems). Control probes detecting a validated positive housekeeping gene (mouse *Ppib* to confirm adequate RNA preservation and detection, 313918, Advanced Cell Diagnostics) and a negative bacterial gene (dapB to confirm absence of nonspecific staining, 312038, Advanced Cell Diagnostics) were used. The chromogen was Fast Red and sections were counterstained with hematoxylin. Positive RNA hybridization was identified as punctate chromogenic red dots under bright field microscopy.

### Adenine Diet Model

#### Experimental Design

Adult female B6 mice (6-8 weeks of age; n=60) were utilized. Mice were randomly assigned by a random number generator (Randomizer.org) to 1 of 2 groups. Thirty mice were inoculated with MKPV and 30 were sham inoculated with an equivalent volume of sterile PBS. Eight to 10 weeks following inoculation, urine was collected individually from all mice and assayed for MKPV by PCR to confirm infection or lack thereof. Beginning 15 weeks after inoculation, mice were fed a purified, casein-based diet containing 0.2% adenine (TD.140290, Envigo, Madison, WI) ad libitum for a maximum of 8 weeks. Twenty-four hours prior to diet initiation (week 0), blood and urine were collected from each animal. Mice were weighed weekly after diet initiation. On weeks 2, 4, and 8 after diet initiation, 10 mice per group were euthanized by CO_2_ asphyxiation and exsanguinated via caudal vena cava puncture. Complete blood count (CBC), comprehensive serum chemistry analysis, and a complete gross necropsy were performed. The cranial pole of the left kidney was frozen at -80°C and retained for MKPV PCR, and the remaining renal tissues were preserved in 10% neutral buffered formalin for histopathology. On weeks 0, 2, 4, 6, and 8 after diet initiation, blood was collected via submental puncture from all surviving mice and serum blood urea nitrogen (BUN), creatinine, and phosphorus were quantified. Additionally, serum SDMA levels were determined in all mice (n=20; 10 per infection status) surviving until the 8-week time point. Two mice in the uninfected group were euthanized after the 6-week time point, but before the 8-week time point, as they reached humane endpoint criteria. Therefore, the group size for the uninfected mice at the 8-week timepoint was 8.

#### Clinical Pathology

Complete blood counts (CBC) were performed using an automated analyzer (ProCyte Dx Hematology Analyzer, IDEXX Laboratories, Inc., Westbrook, ME) within 2 hours of blood collection into collection tubes containing anti-coagulant (MiniCollect K2E K2EDTA). The following parameters were determined: white blood cell count, red blood cell count, hemoglobin concentration, hematocrit, mean corpuscular volume, mean corpuscular hemoglobin, mean corpuscular hemoglobin concentration, red blood cell distribution width standard deviation and coefficient of variance, reticulocyte relative and absolute counts, platelet count, platelet distribution width, mean platelet volume, and relative and absolute counts of neutrophils, lymphocytes, monocytes, eosinophils, and basophils.

For serum chemistry analysis, clotted blood collected in serum separator tubes (MiniCollect 0.8mL CAT Serum Separator, Greiner Bio-One, Monroe, NC) was centrifuged at 3913 x g for 10 minutes, the serum decanted and stored at -80°C until analysis. Thawed serum samples were processed using an automated analyzer (AU680 Chemistry Analyzer, Beckman Coulter, Brea, CA). The concentration of several or all of the following analytes was determined, depending on the experimental aim, as described above: albumin (ALB), total protein (TP), globulin (GLOB), blood urea nitrogen (BUN), creatinine (CREA), calcium (Ca), phosphorus (P). BUN/creatinine and albumin/globulin ratios were calculated. Symmetric dimethylarginine (SDMA) was extracted from 20 uL samples (serum) as well as calibrators and quality control standards, and measured by liquid chromatography-mass spectrometry as previously described (IDEXX Laboratories, Inc., Westbrook, ME)^19^.

Urine collected in preservative-free collection tubes (0.2mL PCR Tube) was processed using the AU680 Chemistry Analyzer to determine urine total protein and creatinine concentration. A urine total protein/creatinine concentration ratio was calculated. Urine specific gravity was measured with a refractometer (TS Meter Refractometer, Leica, Buffalo, NY).

#### Histopathology

Following euthanasia by CO_2_ asphyxiation, a complete post-mortem gross examination was performed, gross lesions were recorded, and all tissues were fixed in 10% neutral buffered formalin. After 24-48 hours of fixation, kidneys were trimmed into two longitudinal and one transverse sections per mouse, processed in alcohol and xylene, paraffin-embedded, sectioned at 5 microns and stained with hematoxylin and eosin (H&E) and the Sirius Red histochemical reaction for collagen.

Renal lesions in the H&E-stained sections were scored using a semi-quantitative method where the following features were assessed and scored: extent of tubular degeneration (estimated as the percentage of tubules with degenerative changes, including epithelial basophilia, vacuolation, attenuation, and necrosis, evaluated at 4x magnification utilizing the scoring scale: 0: 0%; 1: 1-50%; 2: 51-60%; 3: 61-70%; 4:71-80%; 5:81-90%; 6: 91-100%), intra-tubular inflammation (number of tubules with intratubular inflammation in ten 10x fields), interstitial lymphoplasmacytic infiltrates (number of interstitial lymphoplasmacytic infiltrate foci visible at 4x on the 3 whole sections), number of mineralized tubules (number of mineralized tubules visible at 4x on the 3 whole sections), and number of adenine crystals (number of adenine crystals in ten 10x fields).

All scoring was performed by two blinded investigators (AR and SM), using different microscopes (AR: Olympus BX61, SM: Olympus BX65) and the average of the two scores was reported. The extent of fibrosis was quantified on whole slide images of Sirius Red stained sections acquired under polarized light using a dual CCD camera with a 20x/0.8NA objective (Olympus BX61 microscope, DP80 camera, CellSens Dimension [v3.2] software, Tokyo, Japan), resulting in an image resolution of 0.51 μm/pixel. The proportion of renal tissue exhibiting birefringence, excluding the renal pelvis, capsule, and interlobar vessels, was quantified using image analysis software (QuPath v0.3.0 thresholder algorithm, Queen’s University, Belfast, Northern Ireland).^3^

#### Immunohistochemistry

Immunohistochemistry (IHC) was performed on kidney sections using an automated stainer and system reagents (Leica Bond RX, Leica Biosystems, Buffalo Grove, IL). IHC stains were selected based on their routine use in published studies of the adenine diet model and included the F4/80 stain for murine macrophages and the KIM-1 and NGAL stain for renal tubular injury^9, 28, 51, 60, 62, 65^. After deparaffinization and heat induced epitope retrieval in a citrate pH 6.0 buffer (F4/80, KIM-1) or EDTA pH 9.0 buffer (NGAL), IHC for F4/80, KIM-1 and NGAL was performed with primary and secondary antibodies as follows: Invitrogen #14-4801-85 (1:100) & Vector Laboratories #BA-4001 (1:100); R&D Systems # AF1817 & Vector Laboratories #BA-5000 (1:1000); Invitrogen PA5-79590 & Leica Biosystems #DS9800 reagent #3 (used at concentration provided by vendor), respectively, followed by a polymer detection system (DS9800, Leica Biosystems). The chromogen was 3,3 diaminobenzidine tetrachloride (DAB) and sections were counterstained with hematoxylin. Slide image acquisition was performed as described for Sirius Red stained sections, except without polarized light. The proportion of tissue area positive for DAB was quantified using image analysis software (QuPath v0.3.0 thresholder algorithm, Queen’s University, Belfast, Northern Ireland).^3^

### Statistical Analysis

For the pharmacokinetic study, drug concentration at each timepoint and PK parameters (AUC_last_, C0, Cl, t_1/2_) were compared by 2-way ANOVA for the variables “strain” and “infection status”. Clinical and histopathology outcomes were compared by 3-way ANOVA for the variables “drug”, “strain”, and “infection status”. Renal clearance was compared by 3-way ANOVA for the variables “collection interval”, “strain”, and “infection status”. If collection interval as a factor was not significant, it was excluded as a variable and a 2-way ANOVA for “strain” and “infection status” was then performed. Post-hoc analysis by Tukey’s multiple comparison test was performed. For the adenine diet model, body weight, serum BUN, creatinine, phosphorus, SDMA, and all urine chemistry parameters were assessed using a repeated measures mixed-effects model of “infection” and “time post-diet initiation”. Post-hoc analysis by Sidak’s multiple comparison test was performed. Correlation between SDMA and creatinine was performed using Pearson’s r. Data that was only collected at terminal time points, including histopathologic outcomes (except extent of tubular degeneration), renal weight, CBC, and complete serum chemistry were assessed by 2-way ANOVA of the variables “infection” and “time post-diet initiation”. Post-hoc analysis by Tukey’s multiple comparison test was performed. Ordinal data (histopathologic semi-quantitative scoring) was analyzed using the Mann-Whitney U test for both pharmacokinetic and adenine diet aims. All analyses were performed using statistics software (Graph Pad Prism 9.1.0, La Jolla, CA). A P value of less than or equal to 0.05 denoted statistical significance.

### Results Pharmacokinetics

#### MKPV Infection

Urine and renal tissue collected from all sham-inoculated mice (n=48) 10-weeks post-inoculation (PI) and at euthanasia were PCR negative for MKPV. Urine collected from NSG mice 11 weeks PI was positive for MKPV in 14 of 24 (58%) of inoculated animals. In contrast, all (n=24) B6 mice that were inoculated for MKPV had PCR positive urine by 11 weeks PI. Kidneys were assayed for MKPV at 14 weeks PI with 6 additional MKPV-inoculated NSG mice testing positive, resulting in an overall infection rate of 83% (20/24). The 4 MKPV negative NSG mice, consisting of 1 mouse receiving methotrexate and 3 receiving lenalidomide, were excluded from pharmacokinetic and pathologic analysis. MKPV RNA was not detected in PCR negative kidneys using in-situ hybridization.

#### Histopathology

The histopathology scores assigned to renal tissues are provided in Figure 1. Drug administered was not a significant factor as determined by 3-way ANOVA and therefore was not analyzed as a variable. Four MKPV inoculated, but PCR negative NSG mice, were inclusion body free and therefore were not scored for histologic changes. One B6 mouse inoculated with MKPV spontaneously died for reasons unrelated to the study and therefore its kidneys were not assessed. Infected B6 mice had significantly greater interstitial lymphoplasmacytic infiltrates than uninfected B6 mice (*P*<0.0001). Tubular degeneration was significantly more severe in MKPV-infected as compared to sham inoculated NSG (*P*=0.0046) and B6 (*P*=0.0046) mice. While this result was statistically significant, the magnitude of the change was mild; the mean score assigned to infected NSG mice was only 1.1, representing minimal pathology. Eleven of 20 (55%) MKPV positive NSG mice had inclusion bodies in the proximal renal tubular epithelium, whereas only 1 MKPV-positive B6 mouse had inclusion bodies.

**Figure 1:**
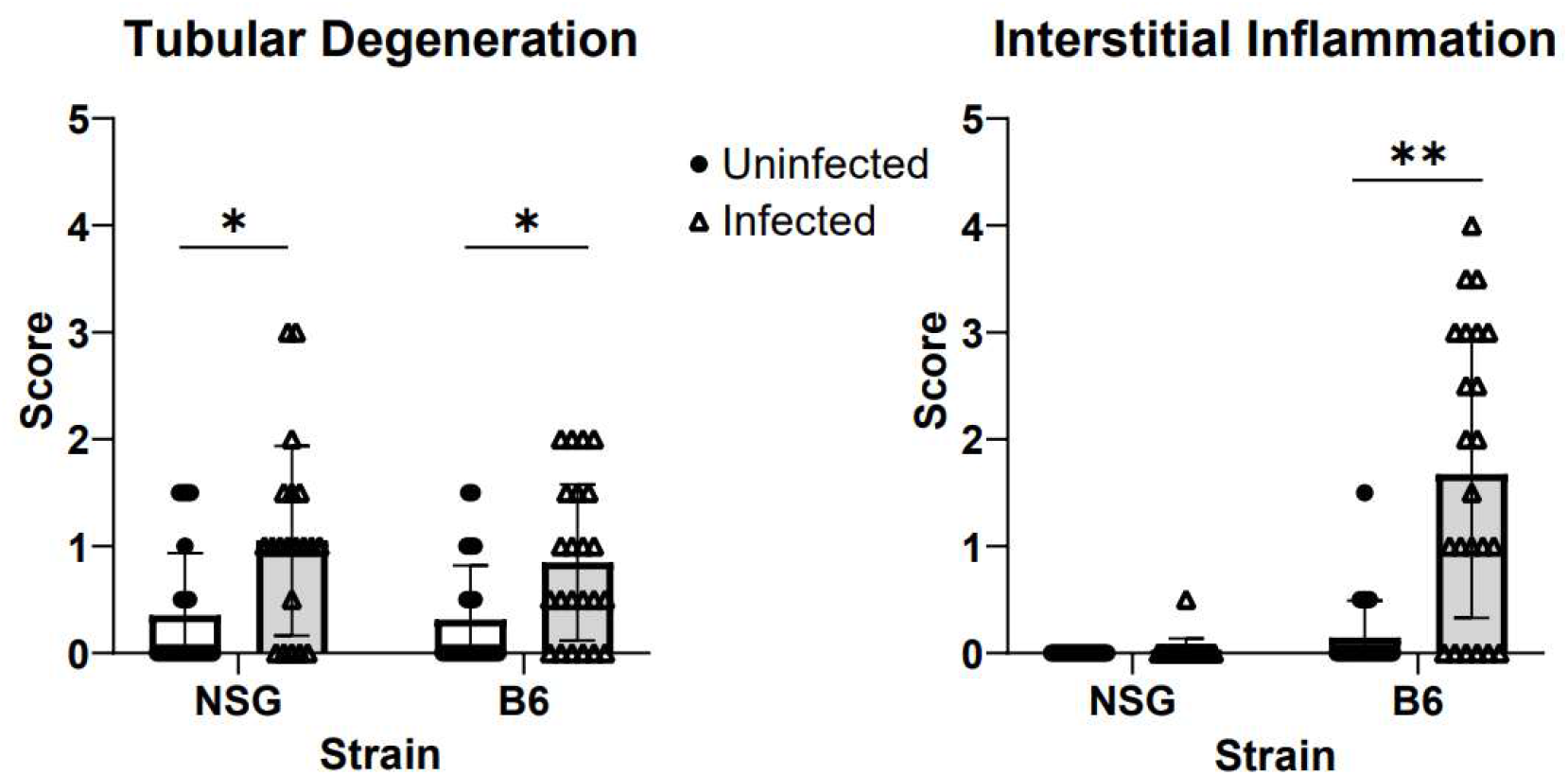
Semi-quantitative scoring of tubular degeneration and interstitial inflammation at 14 weeks PI in NSG and B6 uninfected (closed circle) and infected (open triangle) mice independent of drug administration. Data displayed as mean (bar) ± standard deviation (error bars) with individual animals (symbols). Group size (n=24) except infected NSG mice (n=20) and infected B6 mice (n=23). \**P*<0.005, ** <0.0001.

#### Clinical Pathology

The serum chemistry values from blood samples collected at euthanasia are provided in Table 1. The effect of drug on serum BUN was significant, with mice receiving methotrexate having significantly higher serum BUN than mice receiving lenalidomide ([F (1, 83) = 86.33, P < .0001]). Methotrexate-treated, MKPV-infected NSG mice had significantly higher serum BUN (*P*=0.003) and creatinine (*P*=0.03) than methotrexate-treated, MKPV-infected B6 mice. Methotrexate-treated, MKPV-infected B6 mice had significantly lower serum creatinine than uninfected B6 mice receiving the same drug (*P*= 0.0165). There was no significant impact of strain, drug, or infection status on serum phosphorus concentration.

**Table 1:** Serum chemistry analytes at 14 weeks post-infection in NSG and B6 mice receiving methotrexate or lenalidomide

| Drug | Strain | Infection Status | BUN (mg/dL) | Creatinine (mg/dL) | Phosphorus (mg/dL) |
| --- | --- | --- | --- | --- | --- |
| Methotrexate | NSG | Uninfected | 34.50 ± 5.27<sup>c</sup> | 0.32 ± 0.09 | 11.29 ± 1.52 |
|  |  | Infected | 39.18 ± 4.79<sup>b, c</sup> | 0.33 ± 0.07<sup>b</sup> | 12.43 ± 2.19 |
|  | B6 | Uninfected | 34.17 ± 4.45<sup>c</sup> | 0.33 ± 0.07<sup>a</sup> | 12.98 ± 1.94 |
|  |  | Infected | 31.58 ± 6.35<sup>b, c</sup> | 0.25 ± 0.03<sup>a, b</sup> | 11.29 ± 1.22 |
| Lenalidomide | NSG | Uninfected | 27.25 ± 4.77<sup>c</sup> | 0.30 ± 0.06 | 11.78 ± 1.55 |
|  |  | Infected | 24.33 ± 3.08<sup>c</sup> | 0.26 ± 0.03 | 11.47 ± 1.76 |
|  | B6 | Uninfected | 25.42 ± 4.40<sup>c</sup> | 0.26 ± 0.07 | 11.73 ± 1.68 |
|  |  | Infected | 25.09 ± 3.94<sup>c</sup> | 0.23 ± 0.03 | 11.66 ± 1.34 |

Data are displayed as mean ± SD. Group size (n=12) except infected NSG mice administered methotrexate (n=11), infected B6 mice administered lenalidomide (n=11) and infected NSG mice administered lenalidomide (n=9).<sup>a</sup>P<0.05 between mice of differing infection status, but the same drug and strain. <sup>b</sup>P<0.05 between mice of differing strain, but the same drug and infection status. <sup>c</sup>P<0.05 between mice receiving differing drugs, but the same infection status and strain.

#### Methotrexate pharmacokinetics (PK)

The post-injection plasma concentrations of methotrexate are shown in Figure 2. One NSG mouse inoculated with MKPV, but PCR negative, was excluded from analysis, which caused the exclusion of a data point at 45 and 240 minutes after drug administration for this group. The plasma concentration of methotrexate was significantly higher in uninfected NSG mice as compared to infected NSG mice at 5 (P=0.0006) and 15 (P=0.04) minutes post-injection. The plasma concentration of methotrexate was significantly higher in infected B6 mice compared to uninfected B6 mice at 5 minutes post-injection (P=0.01). The plasma concentration of methotrexate was significantly higher in uninfected NSG compared to uninfected B6 mice at 5 (P<0.0001), 15 (P=0.002), 30 (P<0.0001), 45 (P=0.009), and 60 (P=0.0009) minutes post-injection.

**Figure 2:**
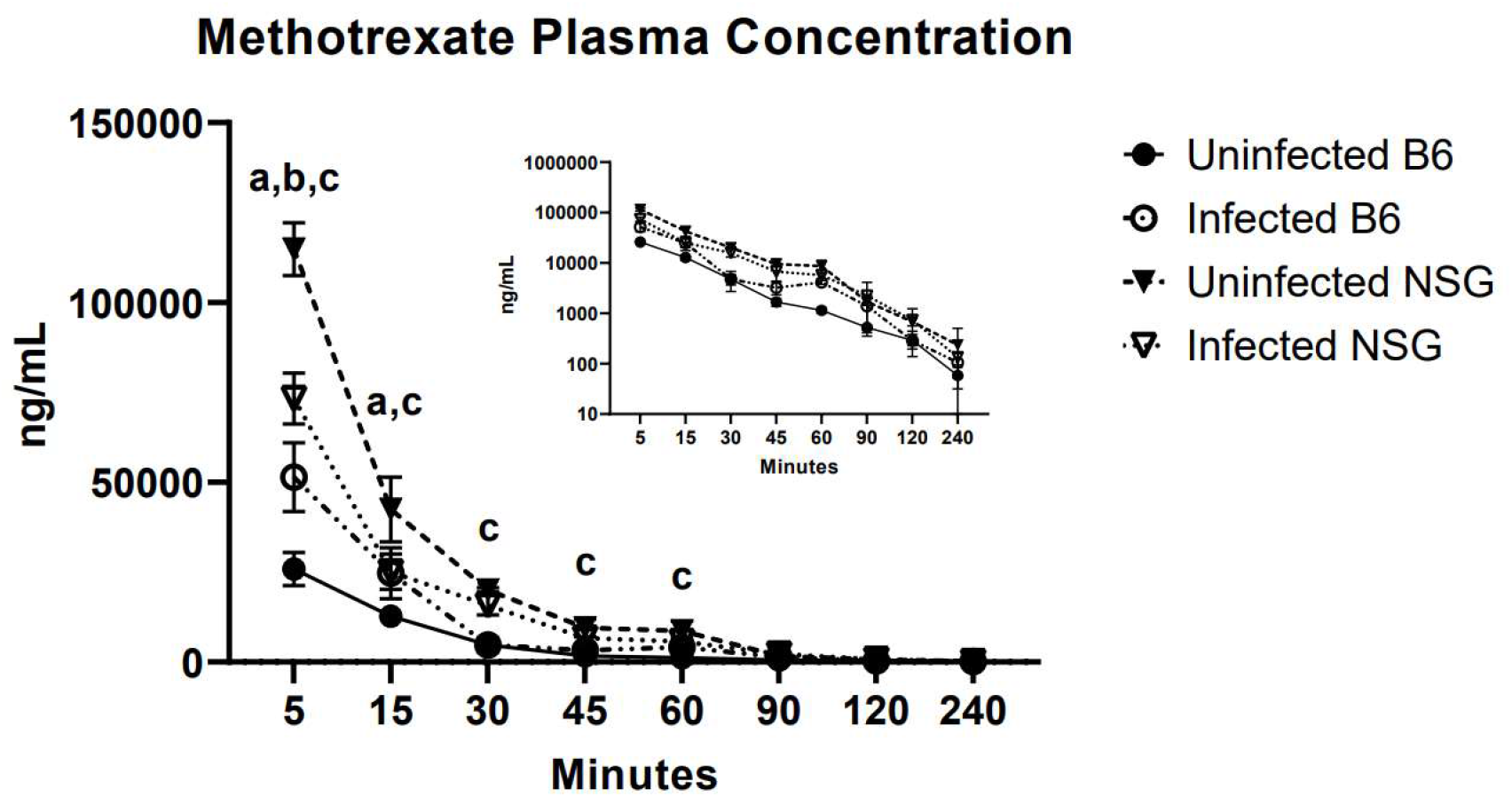
Methotrexate plasma concentrations for uninfected (filled) or infected (open), B6 (circles) or NSG (triangles) mice 14 weeks PI. Log-transformed plasma concentrations in inset. Data are displayed as mean ± SD. Group size (n=3 per time point) except for infected NSG mice at 45 and 240 minutes (n=2). ^a^P<0.05 when comparing infected and uninfected NSG mice. ^b^P<0.05 when comparing infected and uninfected B6 mice. ^c^ = P<0.05 when comparing uninfected NSG and B6 mice.

Noncompartmental (NCA) PK parameters for methotrexate are provided in Table 2. The area under the curve (AUC_last_) was significantly higher in uninfected NSG mice compared to infected NSG mice (P<0.0001), infected B6 mice compared to uninfected B6 mice (P<0.0001), and uninfected NSG mice compared to uninfected B6 mice (P<0.0001). The AUC_last_ was 1.5-times higher in uninfected NSG mice compared to infected NSG mice, 1.9-times higher in infected B6 mice compared to uninfected B6 mice, and 4.3-times higher in uninfected NSG mice compared to uninfected B6 mice. The extrapolated initial concentration (C0) was significantly higher in uninfected NSG mice compared to infected NSG mice (P=0.03) and uninfected B6 mice (P=0.0001). Clearance took significantly longer in uninfected B6 mice as compared to infected B6 mice (P=0.0009) and uninfected NSG mice (P<0.0001). The half-life was not significantly different between strains or infection status.

**Table 2:** Methotrexate non-compartmental pharmacokinetic parameters for uninfected and MKPV-infected B6 and NSG mice 14 weeks post-infection.

|  | B6 |  |  |  |  |  | NSG |  |  |  |  |  |
| --- | --- | --- | --- | --- | --- | --- | --- | --- | --- | --- | --- | --- |
|  | Uninfected |  |  | Infected |  |  | Uninfected |  |  | Infected |  |  |
|  | Geometric Mean | 95% CI | %CV | Geometric Mean | 95% CI | %CV | Geometric Mean | 95% CI | %CV | Geometric Mean | 95% CI | %CV |
| AUC<sub>last</sub> (min*ng/mL) | 603,460<sup>a, b</sup> | 409,423 – 889,456 | 16.25 | 1,169,612<sup>a</sup> | 1,146,500 – 1,193,189 | 0.80 | 2,621,271<sup>a, b</sup> | 2,544,772 – 2,700,069 | 1.20 | 1,778,095<sup>a</sup> | 1,551,174 – 2,038,212 | 5.49 |
| C<sub>0</sub> (ng/mL) | 36,331<sup>b</sup> | 21,300 – 61,970 | 22.30 | 73,839 | 26,777 – 203,617 | 35.23 | 189,968<sup>a, b</sup> | 140,196 – 257,410 | 11.87 | 125,527<sup>a</sup> | 77,726 – 202,725 | 18.23 |
| Cl (mL/min) | 3.10<sup>a, b</sup> | 1.90 – 5.05 | 18.45 | 1.56<sup>a</sup> | 1.31 – 1.86 | 7.03 | 0.70<sup>b</sup> | 0.68 – 0.73 | 1.43 | 1.04 | 0.88 – 1.23 | 6.73 |
| T<sub>1/2</sub> (min) | 43.86 | 25.38 – 75.79 | 22.30 | 40.26 | 19.78 – 81.93 | 30.49 | 34.10 | 25.06 – 46.40 | 12.78 | 28.39 | 8.13 – 99.15 | 52.71 |

AUC<sub>last</sub> - area under the concentration-time curve from time zero to the last observation at 240 minutes after injection; C<sub>0</sub> = back-extrapolated plasma drug concentration at time zero; Cl = systemic clearance; T<sub>1/2</sub> - elimination half-life; CI = confidence interval; CV = coefficient of variation. <sup>a</sup>P<0.05 when comparing infected and uninfected mice of the same strain. <sup>b</sup>P<0.05 when comparing uninfected B6 and NSG mice.

Renal clearance data for mice administered methotrexate are provided in Table 3. Renal clearance was significantly influenced by collection period (F [3, 23] = 3.705, *P*=0.0261). Therefore, renal clearance data was analyzed with collection period as a variable. Mice that had urine collection intervals from 5-60 minutes or 15-90 minutes post-administration often failed to produce adequate urine volumes for methotrexate quantification due to the short time intervals. As there was insufficient data collected at these time points, renal clearance data for these intervals were excluded from analysis. There were no significant differences in renal clearance of methotrexate between infected and uninfected mice of either strain. Mouse strain, but not infection status, was a significant source of variation for methotrexate renal clearance (F [1, 15] = 11.25, *P*=0.0043). Renal clearance, in general, took longer in B6 as compared to NSG mice, however, post-hoc analysis failed to detect statistically significant differences between strains for either specific collection interval.

**Table 3:**
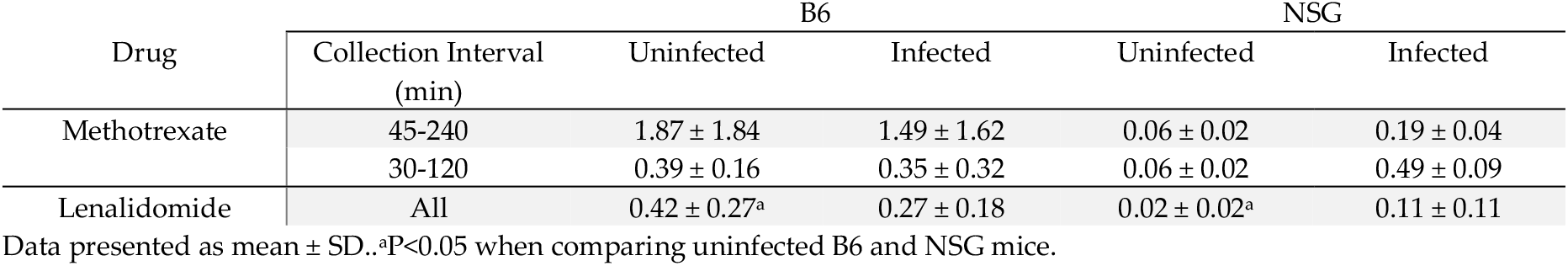
Renal clearance (mL/min) of methotrexate or lenalidomide in uninfected and MKPV-infected B6 and NSG mice 14 weeks post-infection.

| Drug | Collection Interval (min) | B6 |  | NSG |  |
| --- | --- | --- | --- | --- | --- |
|  |  | Uninfected | Infected | Uninfected | Infected |
| Methotrexate | 45-240 | 1.87 ± 1.84 | 1.49 ± 1.62 | 0.06 ± 0.02 | 0.19 ± 0.04 |
|  | 30-120 | 0.39 ± 0.16 | 0.35 ± 0.32 | 0.06 ± 0.02 | 0.49 ± 0.09 |
| Lenalidomide | All | 0.42 ± 0.27<sup>a</sup> | 0.27 ± 0.18 | 0.02 ± 0.02<sup>a</sup> | 0.11 ± 0.11 |

Data presented as mean ± SD. <sup>a</sup>P<0.05 when comparing uninfected B6 and NSG mice.

#### Lenalidomide pharmacokinetics

The post-injection plasma concentrations of lenalidomide are shown in Figure 3. Three inoculated NSG mice were excluded from analysis because they were MKPV PCR negative at the end of the study, which caused the exclusion of a data point at 5, 30, 60, 120, 360, and 480 minutes after drug administration for this group. One MKPV-inoculated B6 mouse was found dead prior to lenalidomide administration and was also excluded, which caused the elimination of a data point at 30 and 360 minutes after drug administration for this group. There were no significant differences in plasma lenalidomide concentrations between infected and uninfected mice at any time point. At 480 minutes post-injection, uninfected NSG mice had a significantly greater lenalidomide plasma concentration than uninfected B6 mice (P=0.0079).

**Figure 3:**
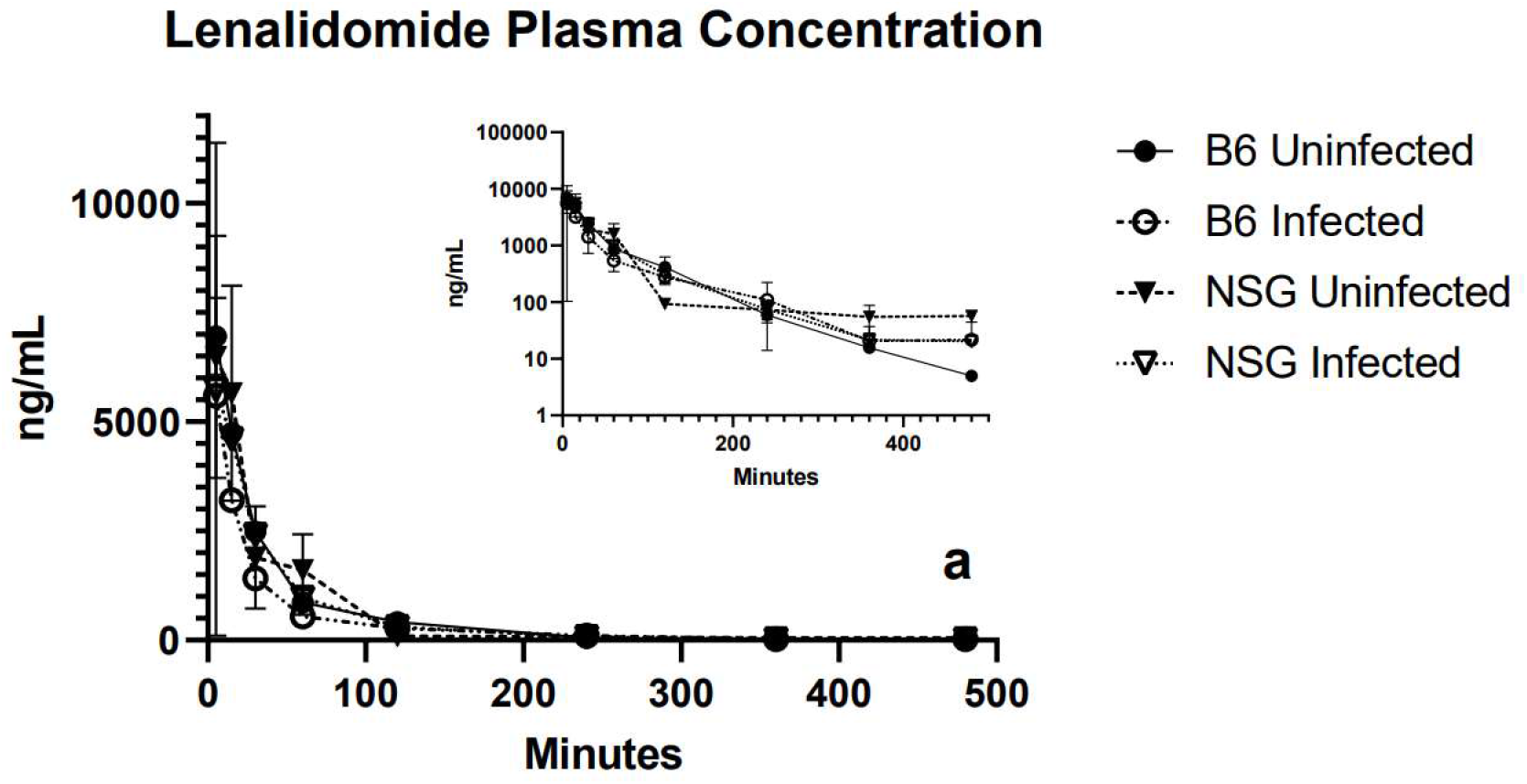
Lenalidomide plasma concentrations for uninfected (filled) or infected (open), B6 (circles) or NSG (triangles) mice 14 weeks PI. Log-transformed plasma concentrations in inset. Data are displayed as mean ± SD. Group size (n=3) except for infected NSG mice at 5, 30, 60, 120, 360, and 480 minutes, and infected B6 mice at 30 and 360 minutes (n=2). ^a^P<0.05 between uninfected NSG and B6 mice.

NCA PK parameters for lenalidomide are provided in Table 4. There were no significant differences in any of the calculated NCA parameters comparing mice of different infection status or strain.

**Table 4:** Lenalidomide non-compartmental pharmacokinetic parameters for uninfected and MKPV-infected B6 and NSG mice 14 weeks post-infection.

|  | B6 |  |  |  |  |  | NSG |  |  |  |  |  |
| --- | --- | --- | --- | --- | --- | --- | --- | --- | --- | --- | --- | --- |
|  | Uninfected |  |  | Infected |  |  | Uninfected |  |  | Infected |  |  |
|  | Geometric Mean | 95% CI | %CV | Geometric Mean | 95% CI | %CV | Geometric Mean | 95% CI | %CV | Geometric Mean | 95% CI | %CV |
| AUC<sub>all</sub> (min*ng/mL) | 273,301 | 220,295 – 339,061 | 8.89 | 209,157 | 160,850 – 271,971 | 10.87 | 282,646 | 240,166 – 331,712 | 6.41 | 290,563 | 191,530 – 440,802 | 16.75 |
| C<sub>0</sub> (ng/mL) | 8,348 | 5,333 – 13,068 | 18.82 | 7,387 | 7,282 – 7,494 | 0.58 | 7,429 | 1,336 – 41,301 | 53.89 | 6,126 | 398 – 94,289 | 73.68 |
| Cl (mL/min) | 0.50 | 0.40 – 0.63 | 8.72 | 0.61 | 0.45 – 0.82 | 11.82 | 0.48 | 0.39 – 0.60 | 8.61 | 0.47 | 0.32 – 0.68 | 14.84 |
| T<sub>1/2</sub> (min) | 41.21 | 33.83 – 50.19 | 8.11 | 75.01 | 53.90 – 104.4 | 12.82 | 69.75 | 55.95 – 86.96 | 8.98 | 64.66 | 22.93 – 182.3 | 44.29 |

AUC<sub>last</sub> - area under the concentration-time curve from time zero to the last observation at 480 minutes after injection; C<sub>0</sub> = back-extrapolated plasma drug concentration at time zero; Cl = systemic clearance; T<sub>1/2</sub> - elimination half-life; CI = confidence interval; and CV = coefficient of variation.

Renal lenalidomide clearance data are provided in Table 3. Renal clearance of lenalidomide was not influenced by time and therefore collection period was not evaluated as a variable. There were no significant differences in lenalidomide renal clearance when comparing infected and uninfected mice of either strain. Strain was a significant source of variation (F [1, 39] = 25.29, *P*<0.0001). Uninfected NSG mice had a significantly reduced renal clearance as compared to uninfected B6 mice (*P*<0.0001).

### Adenine Diet

#### MKPV Infection

Urine and renal tissue collected from all sham-inoculated mice (n=30) 10 wks post-inoculation (PI) and at euthanasia were MKPV PCR negative. Urine collected from all MKPV-inoculated B6 mice (n=30) were MKPV PCR positive by 11 wks PI.

#### Body and Kidney Weight

While both total body weight and renal weight declined after initiating the adenine diet, there were no significant differences in body or renal weight at necropsy at any time point when comparing infected and uninfected mice (Figure 4a; b).

**Figure 4:**
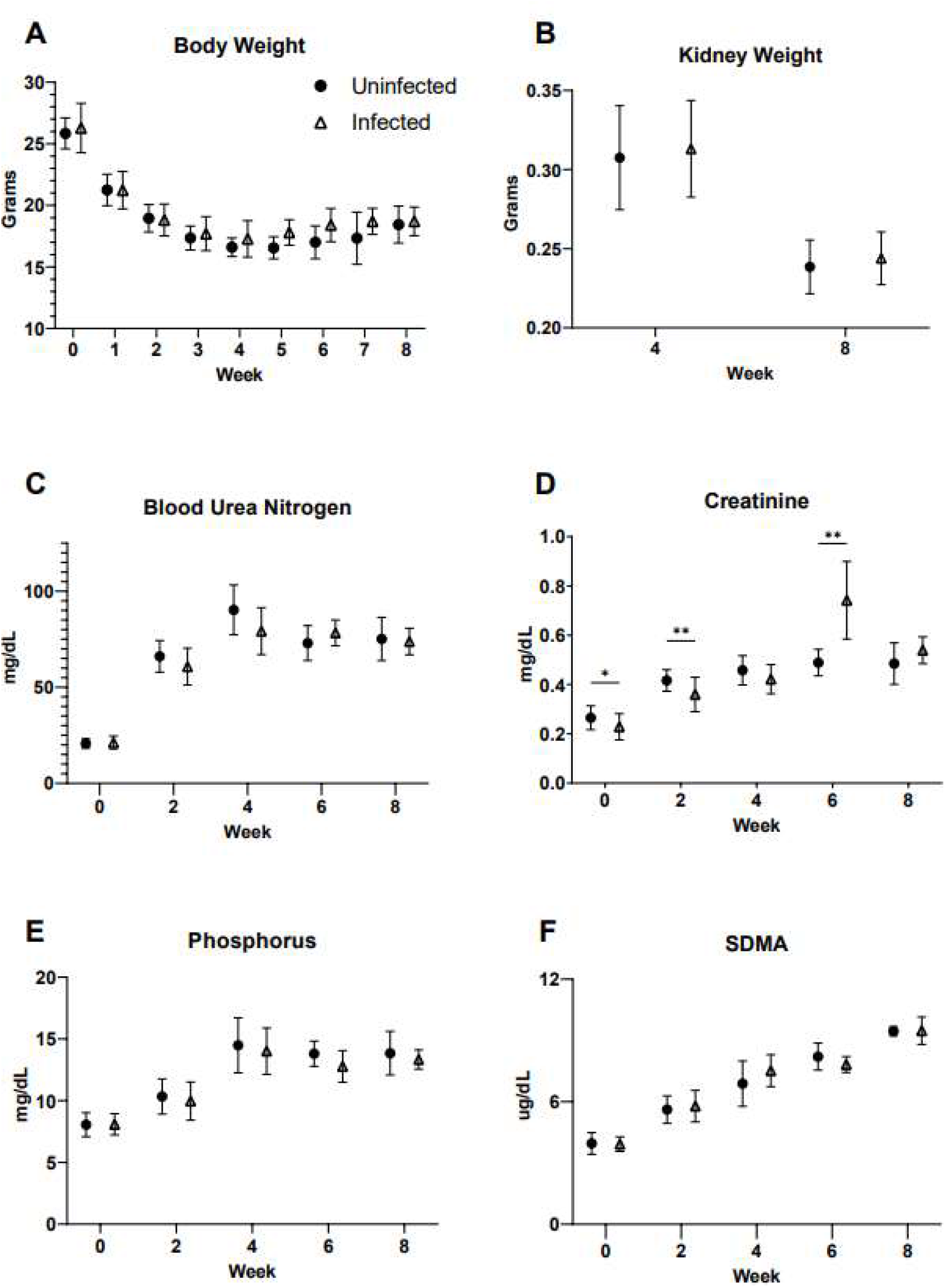
Body and kidney weights and serum renal function analytes in uninfected (circle) and MKPV-infected (triangle) B6 mice after consuming adenine-enriched chow for up to 8 weeks. Mean (± SD) values for a) body weight, b) combined kidney weight at time of necropsy, c) serum blood urea nitrogen, d) serum creatinine, e) serum phosphorus, and f) serum SDMA. * *P* <0.05, *\*\* P* < 0.01. Group size (n=30) for weeks 0, 1, and 2; (n=20) for weeks 3 and 4; and (n=10) for weeks 5, 6, 7, and 8 except for uninfected mice at weeks 7 and 8 (n=8).

#### Clinical Pathology

Serum chemistry analytes are shown in Figure 4c-f and Table 5. Two mice in the uninfected group died between the 6- and 8-week time point, therefore, at week 8 the group size (n=8) was reduced for the uninfected group. Infection status did not have a significant effect on serum BUN, phosphorus, or SDMA at any time point. Serum creatinine was significantly higher in uninfected mice at baseline (*P*=0.034) and at week 2 (*P*=0.0021) compared to infected mice. At 6 wks post-diet initiation, MKPV-infected mice had significantly higher serum creatine than uninfected mice (*P*=0.003), which was not sustained at 8 wks. Other serum chemistry parameters evaluated were not significantly different between infected and uninfected mice at any time point. Creatinine and SDMA were strongly correlated (r=0.68, *P*<0.0001).

**Table 5:** Serum chemistry analytes from uninfected and MKPV-infected B6 mice following 2, 4, or 8 weeks of consuming adenine-enriched chow.

|  | Week 2 |  | Week 4 |  | Week 8 |  |
| --- | --- | --- | --- | --- | --- | --- |
|  | Uninfected | Infected | Uninfected | Infected | Uninfected | Infected |
| BUN (mg/dL) | 71 ± 5 | 67 ± 11 | 94 ± 15 | 86 ± 12 | 75 ± 11 | 74 ± 7 |
| CREA (mg/dL) | 0.40 ± 0.10 | 0.39 ± 0.05 | 0.46 ± 0.05 | 0.42 ± 0.06 | 0.49 ± 0.08 | 0.54 ± 0.05 |
| BUN/CREA Ratio | 180.0 ± 28.4 | 172.6 ± 25.6 | 202.7 ± 15.5 | 205.5 ± 18.5 | 164.9 ± 70.5 | 137.4 ± 11.8 |
| SDMA (μg/dL) | 5.61 ± 0.7 | 5.78 ± 0.8 | 6.88 ± 1.1 | 7.52 ± 0.8 | 9.45 ± 0.2 | 9.48 ± 0.7 |
| TP (g/dL) | 6.1 ± 0.2 | 6.0 ± 0.3 | 6.2 ± 0.5 | 6.0 ± 0.3 | 6.3 ± 0.2 | 6.3 ± 0.2 |
| ALB (g/dL) | 3.2 ± 0.1 | 3.1 ± 0.1 | 3.0 ± 0.2 | 2.9 ± 0.1 | 3.2 ± 0.3 | 3.1 ± 0.1 |
| GLOB (g/dL) | 2.9 ± 0.2 | 2.9 ± 0.3 | 3.2 ± 0.3 | 3.1 ± 0.2 | 3.1 ± 0.2 | 3.2 ± 0.2 |
| A/G Ratio | 1.1 ± 0.1 | 1.1 ± 0.1 | 1.0 ± 0.1 | 1.0 ± 0.1 | 1.0 ± 0.2 | 1.0 ± 0.1 |
| P (mg/dL) | 11.3 ± 1.3 | 11.2 ± 1.6 | 15.6 ± 2.1 | 15.2 ± 1.6 | 13.8 ± 1.8 | 13.3 ± 0.8 |
| Ca (mg/dL) | 11.3 ± 0.4 | 11.2 ± 0.5 | 10.8 ± 0.5 | 10.9 ± 0.3 | 10.9 ± 1.1 | 11.4 ± 0.5 |

Data are shown as mean ± SD.
Group size (n=10) except uninfected mice at week 8 (n=8).

Complete blood counts are provided in Table 6. There were no significant differences in any of the parameters examined when comparing uninfected and infected mice at any time point. Time post-diet administration, but not infection status, was an important source of variation for multiple parameters. The absolute reticulocyte count increased significantly from week 2 to 8 in both uninfected (*P*=0.0001) and infected (*P*=0.0002) mice. Hematocrit decreased significantly from week 2 to 8 in both uninfected (*P*<0.0001) and infected (*P*=0.0009) mice. For both reticulocyte count and hematocrit, infection status had no effect and there was no difference when comparing infected and uninfected mice at any time point.

**Table 6:** Hematologic parameters from uninfected and MKPV-infected B6 mice at 2, 4, or 8 weeks of consuming adenine-enriched chow.

|  | Week 2 |  | Week 4 |  | Week 8 |  |
| --- | --- | --- | --- | --- | --- | --- |
|  | Uninfected | Infected | Uninfected | Infected | Uninfected | Infected |
| RBC (M/uL) | 10.90 ± 2.25 | 10.50 ± 3.67 | 9.30 ± 1.25 | 10.26 ± 0.81 | 8.41 ± 1.64 | 9.25 ± 0.51 |
| HGB (g/dL) | 14.72 ± 3.13 | 14.01 ± 4.87 | 11.21 ± 1.66 | 12.37 ± 1.07 | 9.04 ± 1.64 | 9.99 ± 0.55 |
| HCT (%) | 45.45 ± 10.90 | 46.24 ± 8.58 | 36.86 ± 6.00 | 41.45 ± 3.71 | 30.48 ± 5.29 | 33.82 ± 2.09 |
| MCV (fL) | 44.11 ± 1.89 | 43.0 ± 2.98 | 39.54 ± 1.47 | 40.36 ± 1.12 | 36.51 ± 2.24 | 36.57 ± 0.88 |
| MCH (pg) | 13.47 ± 0.36 | 13.16 ± 0.80 | 12.05 ± 0.32 | 12.04 ± 0.35 | 10.79 ± 0.44 | 10.81 ± 0.27 |
| MCHC (g/dL) | 30.58 ± 1.08 | 30.63 ± 1.35 | 30.48 ± 0.70 | 29.85 ± 0.39 | 29.59 ± 0.74 | 29.54 ± 0.73 |
| RDW-SD (fL) | 29.83 ± 1.96 | 30.9 ± 3.29 | 32.19 ± 3.05 | 32.35 ± 2.74 | 34.91 ± 7.21 | 31.54 ± 1.74 |
| RDW-CV (%) | 27.37 ± 2.67 | 27.8 ± 3.62 | 31.4 ± 2.90 | 31.93 ± 2.74 | 36.23 ± 3.23 | 33.98 ± 1.50 |
| RET# (K/uL) | 372.89 ± 52.87 | 351.53 ± 204.02 | 468.83 ± 97.91 | 503.89 ± 87.40 | 731.50 ± 377.61 | 618.48 ± 85.81 |
| PLT (K/uL) | 1264.3 ± 387.7 | 1051.44 ± 831.56 | 1853 ± 289.63 | 1814.2 ± 334.94 | 1845.50 ± 600.26 | 1713.90 ± 641.54 |
| PDW (fL) | 8.25 ± 0.50 | 8.51 ± 0.71 | 7.66 ± 0.36 | 7.79 ± 0.37 | 8.12 ± 0.46 | 8.08 ± 0.34 |
| MPV (fL) | 6.8 ± 0.8 | 6.83 ± 0.54 | 6.35 ± 0.88 | 6.58 ± 1.04 | 6.22 ± 0.99 | 5.84 ± 0.23 |
| WBC (K/uL) | 9.99 ± 2.01 | 9.53 ± 2.51 | 3.54 ± 0.74 | 3.62 ± 0.76 | 4.86 ± 1.54 | 5.05 ± 1.58 |
| NEUT (K/uL) | 3.51 ± 0.92 | 4.05 ± 1.72 | 2.31 ± 0.74 | 2.22 ± 0.44 | 1.82 ± 0.65 | 1.84 ± 0.75 |
| LYMPH (K/uL) | 5.87 ± 1.72 | 4.97 ± 1.20 | 1.12 ± 0.47 | 1.29 ± 0.46 | 2.63 ± 1.34 | 2.80 ± 0.91 |
| MONO (K/uL) | 0.46 ± 0.16 | 0.39 ± 0.09 | 0.10 ± 0.02 | 0.10 ± 0.05 | 0.35 ± 0.15 | 0.33 ± 0.10 |
| EO (K/uL) | 0.13 ± 0.05 | 0.10 ± 0.06 | 0.01 ± 0.01 | 0.01 ± 0.01 | 0.06 ± 0.04 | 0.06 ± 0.04 |
| BASO (K/uL) | 0.02 ± 0.01 | 0.01 ± 0.01 | 0.01 ± 0.01 | 0.01 ± 0.005 | 0.01 ± 0.01 | 0.01 ± 0.01 |

Data are shown as mean ± SD.
Group size (n=10) except uninfected mice at week 8 (n=8).

#### Urine Chemistry

Urine chemistry analytes are provided in Figure 5. There was no significant difference in urine specific gravity, urinary creatinine, urine total protein, or urine total protein:creatinine ratio when comparing infected and uninfected mice at any time point. All urinary parameters were reduced significantly after adenine diet initiation.

**Figure 5:**
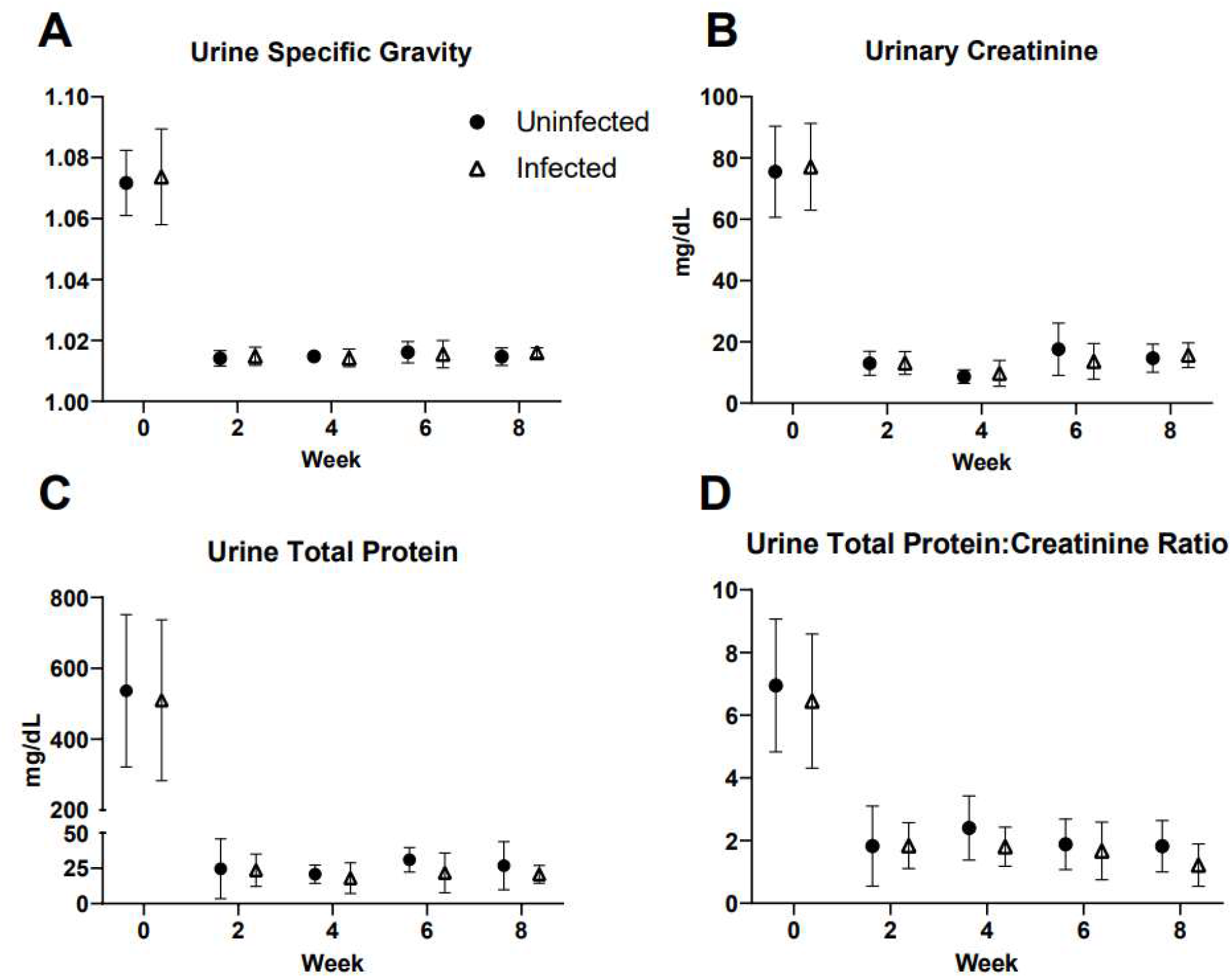
Urine chemistry parameters of uninfected (circle) and MKPV infected (triangle) B6 mice after consuming adenine-enriched chow for up to 8 weeks. Mean (± SD) values for a) urine specific gravity, b) urine creatinine, c) urine total protein, d) urine total protein:creatinine ratio. Group size (n=30) for weeks 0, 1, and 2; (n=20) for weeks 3 and 4; and (n=10) for weeks 5, 6, 7, and 8 except for uninfected mice at weeks 7 and 8 (n=8).

#### Histopathology

##### H&E

Representative photomicrographs of lesions observed after 8 weeks of adenine diet consumption are provided in Figure 6a-d. The scoring of histologic features observed in the adenine diet study is provided in Figure 7. There were no significant differences detected in the extent of tubular degeneration, intra-tubular inflammation, mineralization, or number of adenine crystals between infected and uninfected mice at any time point (Figure 7a-b, d-e). Interstitial lymphoplasmacytic infiltrates were higher in infected as compared to uninfected mice at all time points (Figure 7c). This difference was statistically significant 4-(*P*<0.0001) and 8-(*P*<0.0001) weeks after diet initiation.

**Figure 6:**
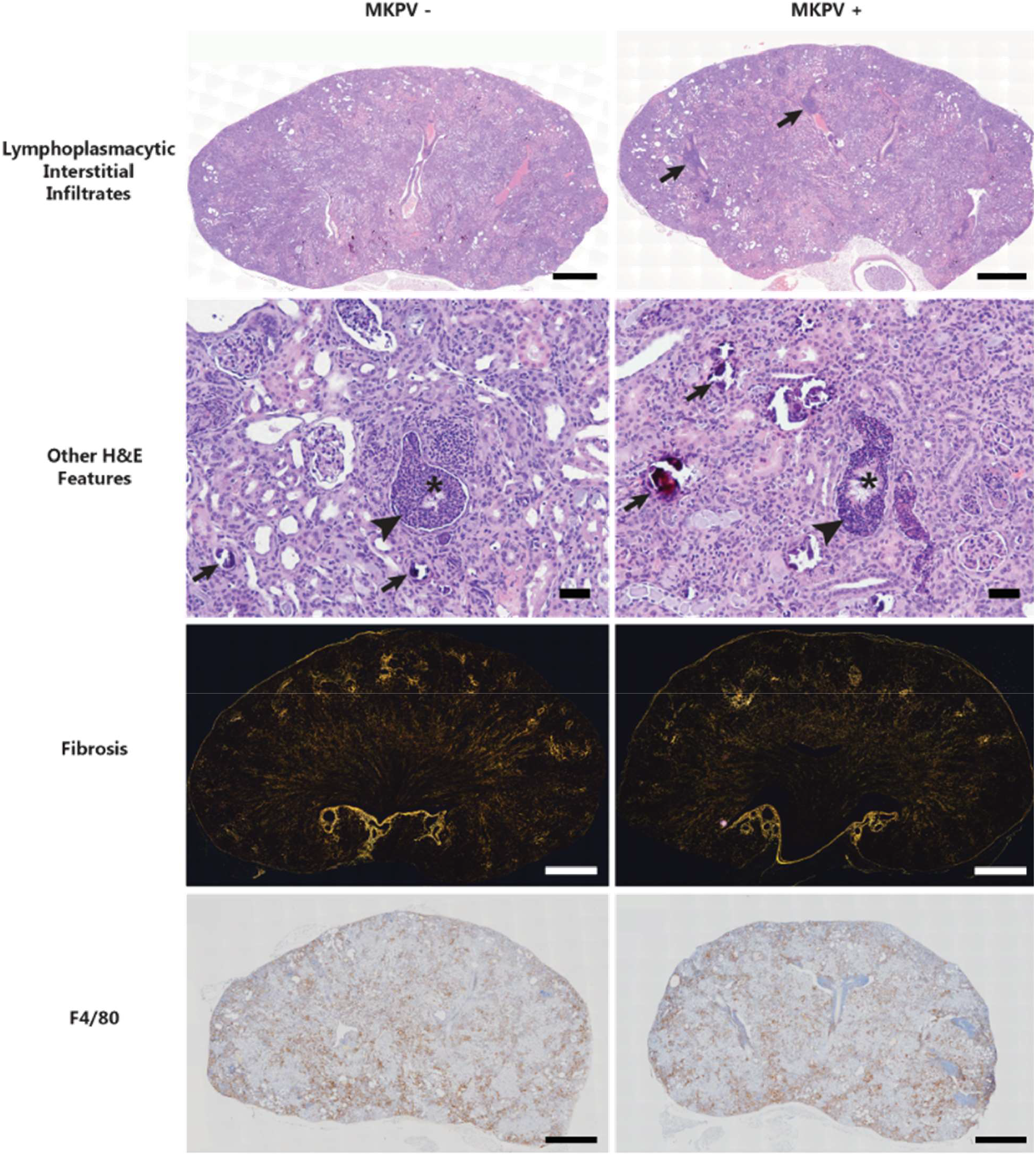
Representative photomicrographs of whole kidney sections from uninfected and MKPV-infected B6 mice following consumption of adenine-enriched chow for 8 weeks. H&E-stained kidney sections demonstrating interstitial lymphoplasmacytic infiltrates (arrows) in uninfected and infected mice. H&E-stained kidney sections demonstrating the remaining features assessed in uninfected and infected mice including mineralization (arrow), intratubular inflammation (arrowhead), and adenine crystals (asterisk). Sirius red-stained kidney sections photographed under polarized light demonstrating interstitial fibrosis in uninfected and infected mice. F4/80 immunohistochemistry kidney sections for uninfected and infected mice. Scale bars: 50 μm (Other H&E Features), 1 mm (all other panels).

**Figure 7:**
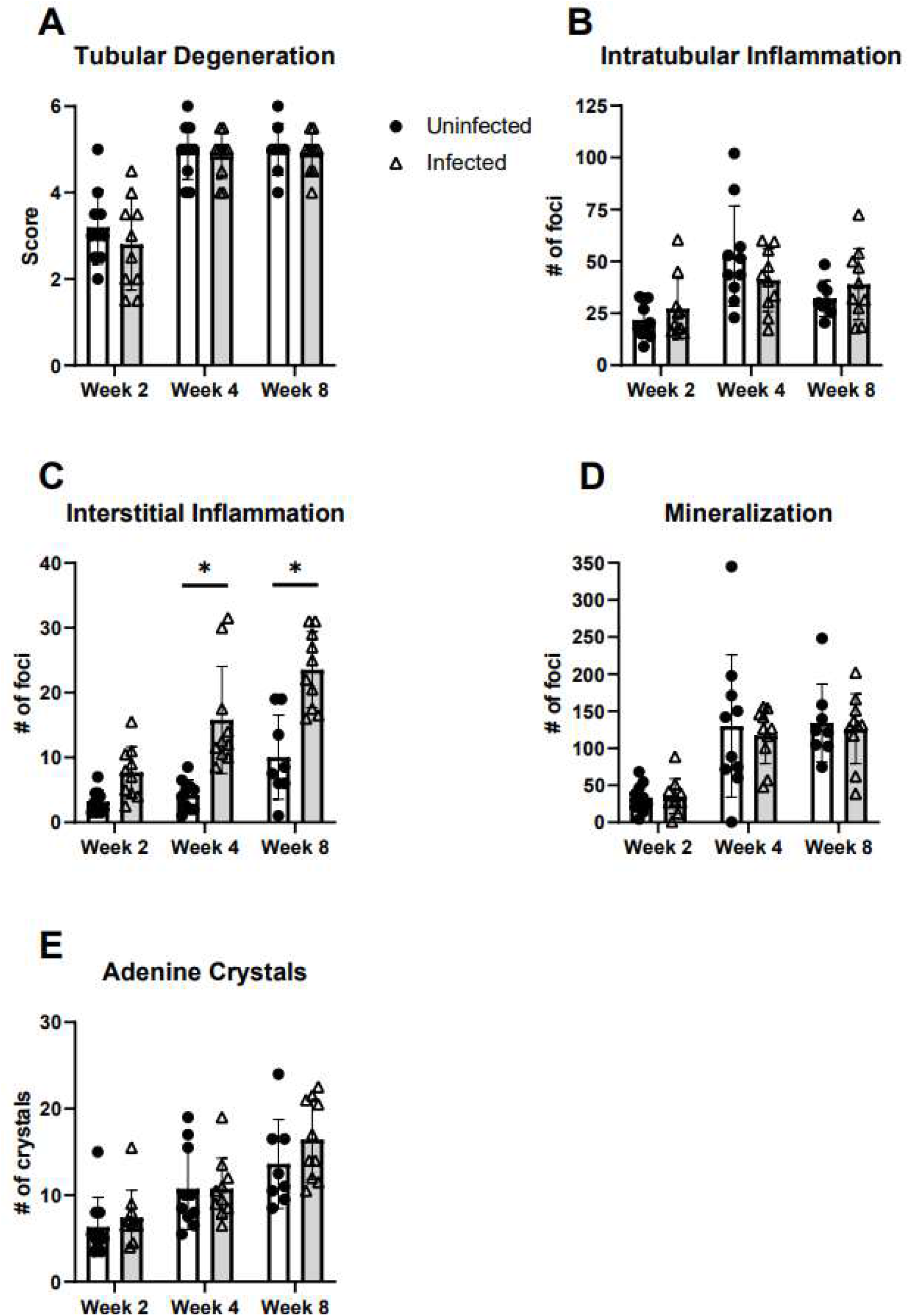
Evaluation of H&E-stained renal tissue in uninfected (circle) and MKPV-infected (triangle) B6 mice following consumption of adenine-enriched chow for 2, 4, or 8 weeks. Data presented as mean ± SD with individual animals (symbols). Group size (n=10) except for uninfected mice at week 8 (n=8). \**P*<0.0001.

##### Sirius Red

Representative photomicrographs of Sirius Red-stained whole kidney sections after 8 weeks of adenine diet consumption are provided in Figure 6e-f. The quantification of renal collagen is provided in Figure 8a. Interstitial fibrosis increased over time. There were no significant differences in the amount of renal fibrosis in infected as compared to uninfected mice 2- and 4-weeks after adenine diet initiation. Infected mice had significantly less interstitial fibrosis at 8-weeks after diet initiation as compared to uninfected mice (P=0.0009).

**Figure 8:**
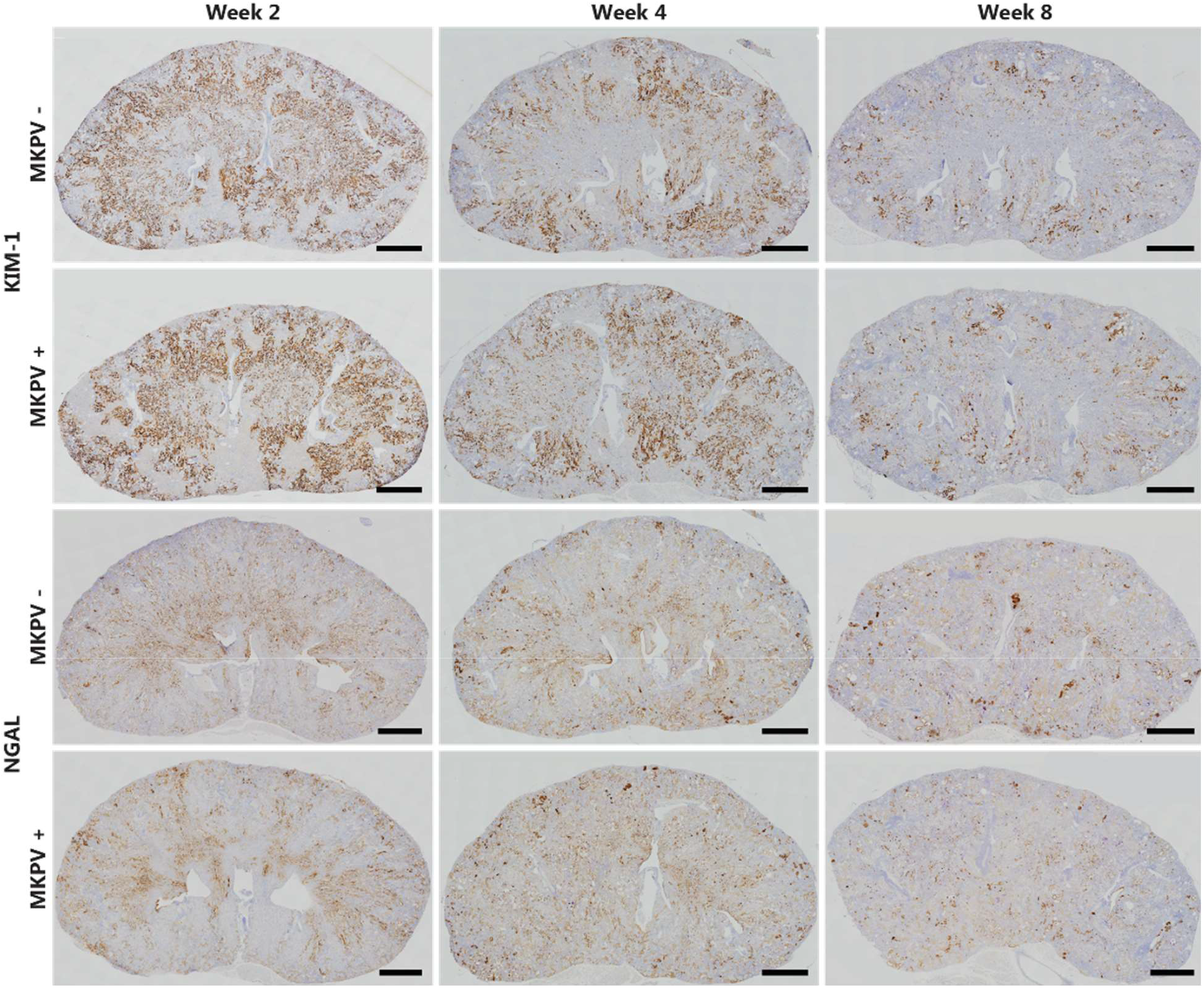
Representative photomicrographs of whole kidney sections from uninfected and MKPV-infected B6 mice following consumption of adenine-enriched chow for 2, 4, and 8 weeks stained by immunohistochemistry for KIM-1 and NGAL. Scale bars: 1 mm.

##### Immunohistochemistry

Representative photomicrographs of immunohistochemically-stained tissue sections are provided in Figure 6g-h and Figure 8. The quantification of the DAB-positive areas as a percentage of the total renal tissue area is provided in Figure 9. There were no significant differences in DAB-positive areas when comparing infected and uninfected mice at any time point for F4/80, KIM-1, or NGAL. Both infected and uninfected mice exhibited significantly less KIM-1 staining over time (F [2, 52] = 117.1, *P* = <0.0001).

**Figure 9:**
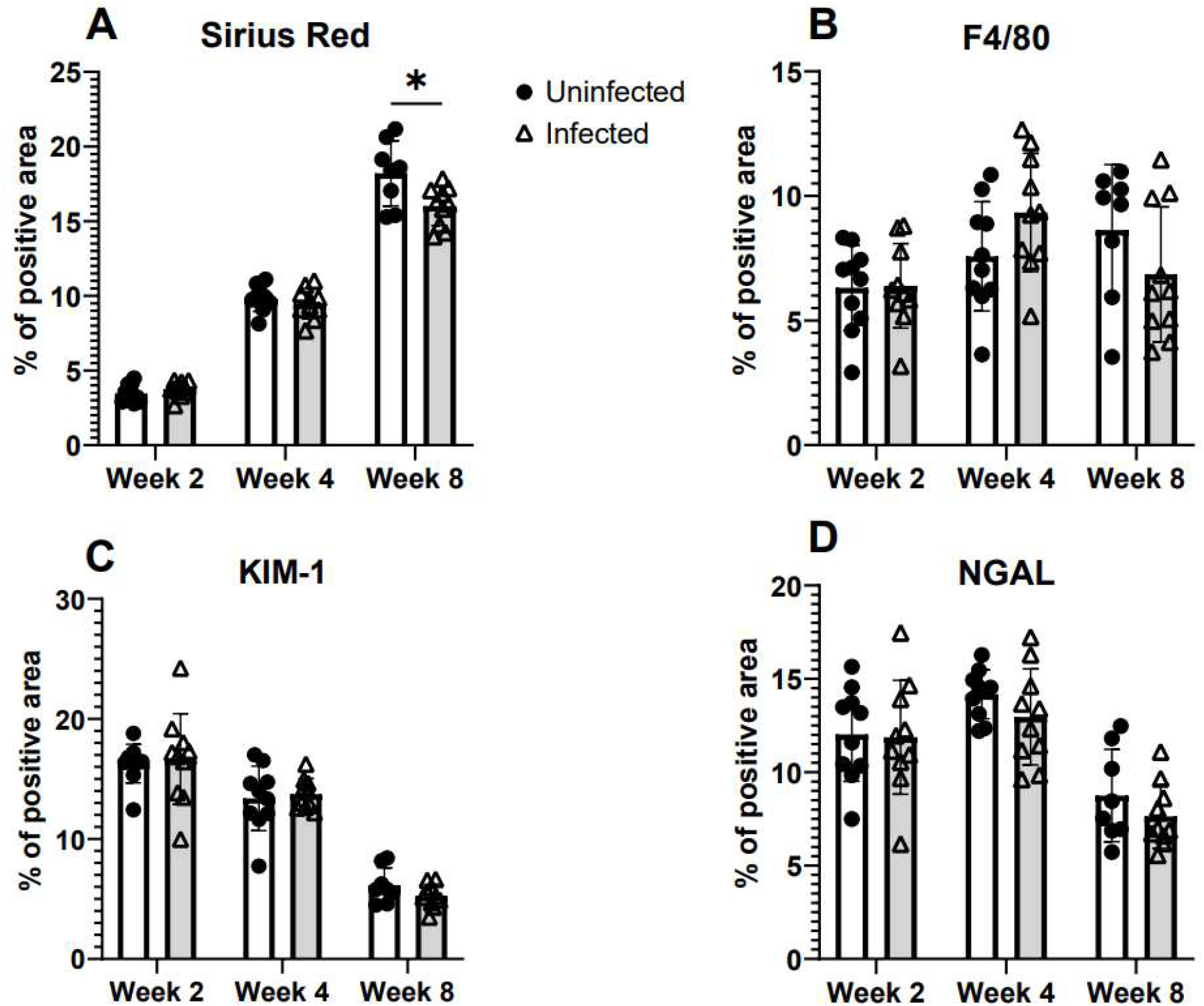
Proportion of renal tissue staining positive for A. Sirius Red, B. F4/80, C. KIM-1 or D. NGAL in naïve (circle) and MKPV-infected (triangle) B6 mice following consumption of adenine-enriched diet for 2, 4, and 8 weeks. Data presented as mean (± SD) with individual animals (symbols) for: a) interstitial fibrosis as determined by percentage of Sirius Red-stained tissue exhibiting birefringence under polarized light, b) F4/80-positive macrophage infiltration as determined by percentage of DAB-positive tissue, tissue injury as determined by percentage of DAB-positive tissue for c) KIM-1 or d) NGAL. \**P* < 0.001. Group size (n=10) except uninfected mice at week 8 (n=8).

## Discussion

All B6 mice inoculated with MKPV were PCR positive by 10-11 wks PI. An appreciable number (4 of 24, 17%) of MKPV inoculated NSG mice did not become infected with the virus, as determined by urine and renal PCR and renal ISH, by 14 wk PI. In our experience with this MKPV infection model, using this inoculation dose and route results in infection detectable by PCR on urine samples in a high proportion of B6 and NSG mice at 10 weeks post-inoculation, and early histopathologic renal changes at 14 to 15 weeks^25^. These results were the basis for the selection of this time point to conduct the PK and initiate the adenine diet studies. However, we have also observed that while the impact of the infection is more severe in NSG than B6 mice after 5 to 6 months, in the early stage of the infection the onset of viral shedding tends occur later and the amount of virus shed in urine tend to be lower in NSG mice as compared to B6^25^. The mechanism(s) for this difference has not been determined. The only sites outside of the kidney where MKPV replication has been documented are in cells within the gastrointestinal lamina propria and in hepatocytes^16; 30^. While viral replication at these sites appeared to be minimal compared to replication in renal tubules, this may play an important role in early infection after oral exposure before the virus reaches the kidney. A possible explanation for the delayed shedding in the early stage of infection in NSG mice is a paucity of the cell(s) that support initial viral replication within the gastrointestinal tract in NSG as compared to B6 mice. These cell(s) may be of immune origin, which may be reduced or lacking in the NSG strain. Murine astrovirus-2 (MuAstV-2) is a recent example of a murine virus that appears to require an immune cell(s) to perpetuate infection as it fails to infect NSG mice^46^.

MKPV infection did not result in significant differences in serum BUN or creatinine concentrations in mice administered chemotherapeutics at 14 weeks PI, except that methotrexate-treated, MKPV-infected B6 mice had significantly lower serum creatinine concentrations as compared to uninfected B6 mice. These results indicate that MKPV infection did not result in overt reduction of kidney function in either strain in this experiment.. All groups of mice administered methotrexate had significantly higher serum BUN concentrations when compared to mice administered lenalidomide, however there was not a concomitant significant increase in creatinine. Elevations in BUN, without a corresponding increase in serum creatinine, can result from proximal gastrointestinal bleeding^17^. Administration of methotrexate has been documented to cause gastrointestinal mucosal injury in humans and mice, however these adverse events are typically associated with chronic administration^12; 63^. In the present study, there was a maximum of four hours between administration of methotrexate and blood collection. Microscopic examination of the gastrointestinal tract was not performed, and therefore the presence of GI mucosal damage was not determined.

There was a significant difference in the AUC_last_ (both strains), clearance (B6 mice only) and C0 (NSG mice only) of methotrexate in MKPV-infected mice when compared to uninfected mice of the same strain. However, there was not a significant difference in half-life observed in either strain, suggesting that changes in the volume of distribution are likely the major contributor to the differences in clearance and C0^36^. Potential mechanisms for a decrease in volume of distribution, which would result in decreased clearance in the MKPV-infected B6 group, would include dehydration or increased drug protein binding. Both of these conditions may occur in disease, but in particular the latter may occur in a pro-inflammatory state associated with systemic infection^37^. MKPV-infected NSG mice had lower C0 than uninfected mice, suggesting a higher volume of distribution. Capillary leakage or decreased drug protein binding are potential causes of increases in volume of distribution. Viral infection can induce endothelial permeability through various cytokine pathways or cause direct endothelial damage^14; 34; 52^. The possibility of vascular permeability contributing to alterations in the volume of distribution in response to infection in the NSG strain is unclear as the *IL2rg*^*null*^ mutation reduces cytokine signaling through multiple receptors.^53; 68^ To the authors knowledge, endothelial damage secondary to MKPV-infection has not been assessed.

Although alterations in AUC between infected and uninfected mice dosed with methotrexate were statistically significant, the biological significance of the differences is less clear. FDA guidance suggests that at least a 2-fold change in AUC is necessary to be considered biologically significant^6; 7; 66^. The difference in AUC between uninfected and infected mice of either strain was less than 2-fold. Additionally, there was no difference in renal clearance of methotrexate between infected and uninfected mice. Renal clearance is typically considered a constant rate when drugs undergo linear kinetics, however methotrexate excretion depends on saturable resorption and secretion pathways, which may explain why clearance was significantly impacted by the collection interval^18^. While there were statistically significant differences in the AUC_last_ and clearance or C0 of methotrexate between infected and uninfected mice, alterations in renal clearance caused by MKPV was likely not the etiology, which was contrary to our initial hypothesis. Without examining pharmacodynamics, one cannot definitively determine whether the alterations in AUC are biologically significant. Differences in methotrexate susceptible tumor growth kinetics could be examined to determine biologic relevance.

There were no significant differences in lenalidomide plasma pharmacokinetics between infected and uninfected mice of the same strain, suggesting that alterations in AUC, clearance, and C0 in infected mice are drug dependent. In humans, there were no significant differences in lenalidomide AUC, half-life, and renal clearance in patients with mild renal impairment as compared to patients with normal renal function, however they were observed in patients with moderate and severe renal impairment^10^. Based on the clinical and anatomic pathology findings in this study, the mice evaluated may reflect early or no functional renal impairment. Although methotrexate and lenalidomide have different mechanisms of renal excretion, the renal clearance of both drugs was not significantly different in MKPV-infected mice as compared to uninfected controls. This suggests that MKPV infection did not significantly impact either renal clearance mechanism. The impact of MKPV infection on other factors that influence plasma pharmacokinetics, including plasma protein binding, drug metabolism, and non-renal excretion, which are different between methotrexate and lenalidomide may have contributed to the difference in impact that MKPV infection had on plasma pharmacokinetics and is an area where additional studies are needed.

Interestingly, we observed significant inter-strain differences in drug disposition, particularly among mice administered methotrexate. Uninfected NSG and B6 mice receiving methotrexate had significant differences in AUC_last_, C0, and clearance, although half-life was not significantly different. However, the magnitude of the difference in half life was larger between uninfected B6 and NSG mice than between infected and uninfected mice of either strain. Significant changes in the volume of distribution along with changes in AUC or clearance may mask alterations in the half-life^5^. Renal clearance of lenalidomide, while not significantly impacted by infection status, was significantly impacted by strain. Possible explanations for these inter-strain pharmacokinetic differences may include differences in plasma protein concentrations or drug binding which can impact volume of distribution and renal clearance of drugs^54^. A study comparing plasma protein profiles of five mouse strains found similar plasma protein profiles across all sexes and strains, including the C57BL/6 and the NOD/SCID mouse^39^. While the plasma protein profile of the NSG mouse was not assessed, the NSG mouse is closely related to the NOD/SCID mouse, differing only by the IL2rg mutation^53^. Therefore, it is likely that the B6 and NSG mouse have similar plasma protein profiles in health; how or if infection impacts the plasma protein profiles of these strains is not documented. While the mechanism(s) underlying the difference in pharmacokinetics is unclear, the differences in pharmacokinetic parameters between strains suggests that pharmacokinetic data should not be extrapolated readily between mouse strains. NSG mice have been demonstrated to have significant differences in the PK profile of antibody and antibody-drug conjugates when compared to BALB/c, athymic nude, and SCID mice, which resulted in reduced antitumor activity of the tested compounds^31^. While the mechanism responsible for the alterations in pharmacokinetic properties of antibodies is likely different than the mechanisms underlying the results in the present study, these results highlight that differences in mouse strain can significantly impact metabolism and excretion of test agents. Therefore, studies investigating pharmacokinetic or toxicokinetic properties of various compounds should be completed in mice of the same strain.

There were several limitations to the pharmacokinetic study. Unfortunately, a few NSG mice needed to be removed from analysis as they did not become infected. Therefore, there were select time points where a smaller number of animals contributed to the PK data, particularly for NSG mice administered lenalidomide, which contributed to greater intra-group variability (%CV). Additionally, we did not assess pharmacodynamic endpoints that would have enhanced our understanding of the biological significance of the alterations in pharmacokinetic parameters. The severity of MKPV infection, based on the histologic changes observed, was relatively mild in the present study. Mice that are at a later stage of infection or have more severe pathology (as commonly occur in older NSG mice after prolonged infection), may have alterations in pharmacokinetics that could not be appreciated. Only female mice were used in this study as prior studies in which MKPV pathogenesis was evaluated temporally was limited to studying female mice^25^. The applicability of this data to other strains of MKPV, later stages of MKPV infection, other strains and sexes of mice, and other drugs requires further study.

In the adenine diet induced model of chronic kidney disease, all inoculated mice became infected prior to induction of the model, and induction of the model was successful in both groups, with infected and uninfected mice developing significant azotemia as well as hyperphosphatemia, anemia, and hyposthenuria relative to baseline, as previously reported in other studies using this model^23; 28; 59^. MKPV infection for >15 weeks did not significantly worsen measurable clinicopathologic parameters in female C57BL/6NCrl mice, which was contrary to our hypothesis that infection would cause significant elevation of renal functional biomarkers in this model. There were no significant differences in serum BUN, phosphorus or SDMA between infected and uninfected mice at any time point. Creatinine was significantly higher in uninfected mice at two timepoints, however the absolute difference between groups was relatively small. At 6-weeks after diet administration, infected mice had significantly higher creatinine than uninfected mice. There were no corresponding elevations in serum SDMA or BUN for infected mice at this time point that would suggest a decreased glomerular filtration rate. Serum samples from infected and uninfected mice were processed for serum creatinine along with BUN and phosphorus in the same analyzer run. Due to concerns regarding the serum creatinine results, analysis was repeated on the same serum samples. These results were not significantly different from the initial results. Based on these factors, it is unlikely that machine error contributed to the significant difference between infected and uninfected groups at this time point. Creatinine was quantified using the Jaffe method, which can be impacted by hemolysis, icterus, lipemia, or binding of non-creatinine chromogens^13; 55; 57^. Due to the repeatability of the elevated creatinine, it is possible that one of these factors could have affected the 6-week timepoint blood samples from the infected group. Therefore, we considered the significant difference in creatinine, when compared to uninfected mice at 6 weeks, to be artifactual.

There was no difference in urine specific gravity, urinary creatinine, urinary total protein, or the urine total protein:creatinine ratio between infected and uninfected mice. Both infected and uninfected mice developed anemia over time, however the nature of this anemia appeared to be regenerative as both groups developed significant increases in absolute reticulocyte count over time in conjunction with significant decreases in hematocrit. This is contrary to the non-regenerative anemia that typically develops in naturally occurring CKD^32^. Previous reports of the adenine diet model have described both non-regenerative anemia as well as anemia with increased absolute reticulocytes as seen in the present study^1; 2; 60^. The mechanism by which these mice develop a regenerative response despite profound renal injury is unclear, but this study adds to the body of evidence that suggests there is variability in the type of anemia that develops in this model.

We were unable to determine if serum SDMA was a more sensitive indicator of renal impairment in the adenine diet model. In other species, as renal disease progresses, SDMA is noted to increase before serum creatinine and is therefore a more sensitive indicator of a decrease in GFR^19; 40^. However, we were unable to observe this trend due to rapid progression of the model. At the earliest time point, 2 weeks after diet initiation, serum creatinine had already increased above the 95% percentile of the baseline serum creatinine in both groups. This rapid progression of azotemia and weight loss appears more severe in the present study than in previous reports of the adenine diet ^11; 23^. A study evaluating earlier time points would be necessary to determine if SDMA increases significantly prior to creatinine. Nevertheless, we did observe that SDMA increased significantly from baseline after initiating the adenine diet and was strongly correlated with serum creatinine. Prior measurement of baseline SDMA concentrations were notably higher (mean 8.4 μg/dL) in male ICR mice as compared to the concentrations in the female C57BL/6NCrl used in the present study (mean 3.9 μg/dL); however, these differences may simply be reflective of differences in strain, sex, and/or analytical method^8^. Additional characterization of baseline SDMA measurements in multiple strains of mice of both sexes is necessary.

Infection with MKPV causes lymphoplasmacytic interstitial inflammation in immunocompetent mice^16^. This was appreciable in both the pharmacokinetic aim, where MKPV-infected B6 mice had significantly higher scores for interstitial inflammation compared to uninfected B6 mice regardless of drug administration, and in the adenine diet aim, where infected mice had significantly greater numbers of lymphoplasmacytic inflammatory foci than uninfected mice at multiple time points. We hypothesized that MKPV-infected mice would develop greater interstitial fibrosis compared to uninfected mice because infection with MKPV has been associated with marked interstitial fibrosis in immunodeficient mice. The limited assessments of MKPV infection in immunocompetent mice did not reveal marked renal fibrosis, but the chronic inflammation that occurs in the kidneys of these mice may promote a milder form of renal fibrosis that could only be documented by special stains, which were not performed in these studies^38; 47^. However, we found a significant reduction of interstitial fibrosis at 8 weeks after adenine diet initiation in mice infected with MKPV as compared to the uninfected mice. Some cytokines that are known to be produced in response to viral infections, including IL-10 and interferon-*γ* (IFNγ), have been associated with downregulation of renal fibrosis. IL-10, a multifunctional cytokine that balances the immune response by acting as a brake on inflammation, is produced by multiple types of immune cells during the acute and chronic stages of viral infections^48^. It has been shown to act as a downregulator of the fibrotic response in multiple organs including the kidney, based on evidence in experimental animal models of chronic kidney disease^56^. Additionally, there is evidence that the pro-inflammatory cytokine IFN*γ*, and a key component of the antiviral immune response, inhibits renal fibrosis in experimental models ^26; 42; 44^. If IFN*γ* or Il-10 were elevated following MKPV infection and immune cell recruitment in immunocompetent mice, these cytokines may have contributed to the attenuation of interstitial renal fibrosis observed in infected mice. Additional studies would be required to determine the effects of MKPV infection on cytokine production in the kidneys of immunocompetent mice and how these effects may modulate renal fibrosis.

Immunostaining for F4/80, a macrophage marker commonly utilized to characterize the adenine diet model, was not significantly different between infected and uninfected mice at any time point. Infection with MKPV has been demonstrated to increase activated renal macrophages in immunodeficient mice^47^. This may not have been appreciated either due to the immunocompetency of the B6 mouse or, if there was a subtle increase in activated renal macrophages, it may have been obscured by the robust macrophage recruitment that occurs in the adenine diet model. KIM-1 (Kidney injury marker 1) staining, a marker of renal tubular damage, which has also been used to characterize renal disease in the adenine diet model, was not significantly different between infected and uninfected mice at any timepoint assessed^28; 51; 62^. However, staining did significantly decline over time, with kidneys collected after 2 weeks of adenine diet consumption demonstrating significantly more staining than kidneys collected after 8 weeks of adenine diet consumption. This is consistent with the other studies that demonstrated that KIM-1 mRNA expression decreased between day 7 and day 14 of adenine diet consumption in wildtype C57BL/6J mice^15^. KIM-1 expression is model-dependent in murine models of CKD, with the unilateral ureteral obstruction model displaying gradually increasing KIM-1 expression over time, while the cisplatin model demonstrated acute KIM-1 expression that resolved by day 14 post-drug administration^61^. We observed that KIM-1 expression is higher following acute injury and declined over time. This may reflect temporally decreasing mass of renal tubular epithelium secondary to tubular injury. The reduction of kidney weight between weeks 4 and 8 of adenine diet consumption supports this observation. Similarly, neutrophil gelatinase-associated lipocalin (NGAL) staining, a marker of renal tubular injury in mice, was not significantly different between infected and uninfected mice at any timepoint^9; 65^. NGAL expression also declined between 2 and 8 weeks after diet initiation; however, it appeared to reach peak expression at week 4, later than KIM-1. NGAL expression is upregulated in injured tubular epithelial cells, but it is also expressed as a component of secondary granules in neutrophils, which may have contributed to increased NGAL staining at this timepoint, as this was also the timepoint where intratubular neutrophilic inflammation peaked^27^.

Although the results of the present study indicate that 15 weeks of infection with MKPV did not significantly exacerbate the measurable clinicopathologic outcomes in the 0.2% adenine diet model of CKD, it did significantly alter select histopathologic features. However, these results cannot necessarily be extrapolated to other models of kidney disease, mouse strains, or stages of infection with the MKPV virus.

The results of this study disproves our initial hypotheses that MKPV infection would significantly alter the renal clearance of the chemotherapeutic drugs methotrexate or lenalidomide and would alter traditional serum and urinary biomarkers of renal function in the adenine diet model of CKD. Together, these results suggest that renal function is not significantly impacted during subclinical MKPV infection. However, infection with MKPV enhanced interstitial lymphoplasmacytic infiltrates and attenuated renal fibrosis at 8 weeks after adenine diet initiation. Based on the present results, if histologic assessment of the kidney is performed in a study utilizing the adenine diet model, MKPV infection status should be documented and reported in order to enhance research reproducibility and allow comparison of results between studies. These results suggest that MKPV may also confound experimental results in other models in which renal histopathology is assessed, but further studies are required to better characterize the impact of MKPV on other models. We did observe mouse strain-specific differences in the pharmacokinetics of the chemotherapeutics evaluated independently of MKPV infection. Therefore, data gathered from pharmacokinetic studies should not be extrapolated between mouse strains without validation.

## Abbreviations and Acronyms

MKPV: Mouse Kidney Parvovirus
IBN: Inclusion body nephropathy
RTE: Renal tubular epithelium
PDX: Patient-derived xenograft
MTX: Methotrexate
LEN: Lenalidomide
DHA: 2,8-dihydroxyadenine
SDMA: Symmetric dimethylarginine
NGAL: Neutrophil gelatinase-associated lipocalin
KIM-1: Kidney injury molecule-1
ISH: In situ hybridization

## Acknowledgements

This work was supported in part by the ACLAM Foundation and the MSK’s NCI Cancer Center Support Grant P30 CA008748. We also acknowledge the contributions of Antonio Bravo, John D’Allara, Jacqueline Candelier, Kim McBride, Maria Jiao, and Iveta Simanska of the Laboratory of Comparative Pathology, Mary Leissinger previously with IDEXX Bioanalytics, and MSK’s Antitumor Assessment Core Facility and Dr. Greg Gorman of Samford University with their assistance with the pharmacokinetics study.

